# Single cell transcriptomics of the *Drosophila* embryonic salivary gland reveals not only induction but also exclusion of expression as key morphogenetic control steps

**DOI:** 10.1101/2024.05.09.593329

**Authors:** Annabel May, Katja Röper

## Abstract

How tissue shape and therefore function is encoded by the genome remains in many cases unresolved. The tubes of the salivary glands in the *Drosophila* embryo start from simple epithelial placodes, specified through the homeotic factors Scr/Hth/Exd. Previous work indicated that early morphogenetic changes are prepatterned by transcriptional changes, but an exhaustive transcriptional blueprint driving physical changes was lacking. We performed single-cell-RNAseq-analysis of FACS-isolated early placodal cells, making up less than 0.4% of cells within the embryo. Differential expression analysis in comparison to epidermal cells analysed in parallel generated a repertoire of genes highly upregulated within placodal cells prior to morphogenetic changes. Furthermore, clustering and pseudo-time analysis of single-cell-sequencing data identified dynamic expression changes along the morphogenetic timeline. Our dataset provides a comprehensive resource for future studies of a simple but highly conserved morphogenetic process of tube morphogenesis. Unexpectedly, we identified a subset of genes that, although initially expressed in the very early placode, then became selectively excluded from the placode but not the surrounding epidermis, including *hth*, *grainyhead* and *tollo/toll-8*. We show that maintaining *tollo* expression severely compromised the tube morphogenesis. *tollo* is likely switched off to not interfere with key Tolls/LRRs that are expressed and function in the placode.

## Introduction

During embryonic development complex shapes arise from simple precursor structures. How organ shape and hence function is encoded by the genome remains in many cases an open question. Although we understand often in great detail how gene regulatory networks are specifying overall organ identity, how such patterning is then turned into physical morphogenetic changes that actually shape tissues is much less understood. We use the formation of the tubes of the *Drosophila* embryonic salivary glands as a simple model of tube morphogenesis through budding, a common pathway to form tubular organs (Iruela-Arispe & Beitel, 2013). The salivary glands are initially specified at stage 10 of embryogenesis as two flat epithelial placodes of approximately 100 cells each on the ventral side of the embryo. The morphogenesis begins with cells in the dorsal-posterior corner constricting their apices and beginning to internalise. Whilst cells disappear through the invagination point on the surface, a narrow lumen tube forms on the inside (Girdler & Röper, 2014; Sanchez-Corrales *et al*, 2018; Sanchez-Corrales *et al*, 2021; Sidor & Röper, 2016).

Studies over the last 30 years have revealed the transcriptional patterning that leads to the specification of the salivary gland placodes, with the key activator being the homeotic transcription factor Sex combs reduced (Scr; (Henderson & Andrew, 2000; Henderson *et al*, 1999)). Scr becomes restricted to parasegment 2 in the embryo through the combined action of homeotic transcription factors T-shirt and Abdominal-B repressing *Scr*’s expression posteriorly (Andrew *et al*, 1994). Dpp signalling affects the dorsal expansion of the Scr domain (Andrew *et al*., 1994; Henderson & Andrew, 2000). Furthermore, classical studies of mutants have revealed many key factors involved in salivary gland morphogenesis (Abrams *et al*, 2003; Sidor & Röper, 2016). The initial primordium, once specified, is quickly subdivided into two groups of cells, secretory cells that will form the main body of the tube, and duct cells, close to the ventral midline, that will eventually form a Y-shaped duct connecting both glands to the mouth, once all secretory cells have internalised (Fig. 1A and Suppl. Fig.1A). These two groups of cells, we know, are established through EGF signalling emanating from the ventral midline and inhibiting Forkhead (Fkh) transcription factor function in the future duct cells. Fkh in the rest of the placodal cells instructs the morphogenesis and activates a secretory programme (Haberman *et al*, 2003; Jones *et al*, 1998). EGFR mutants do not form a duct and internalise salivary gland tubes with two closed ends (Kuo *et al*, 1996; Maybeck & Röper, 2009).

**Figure 1.**
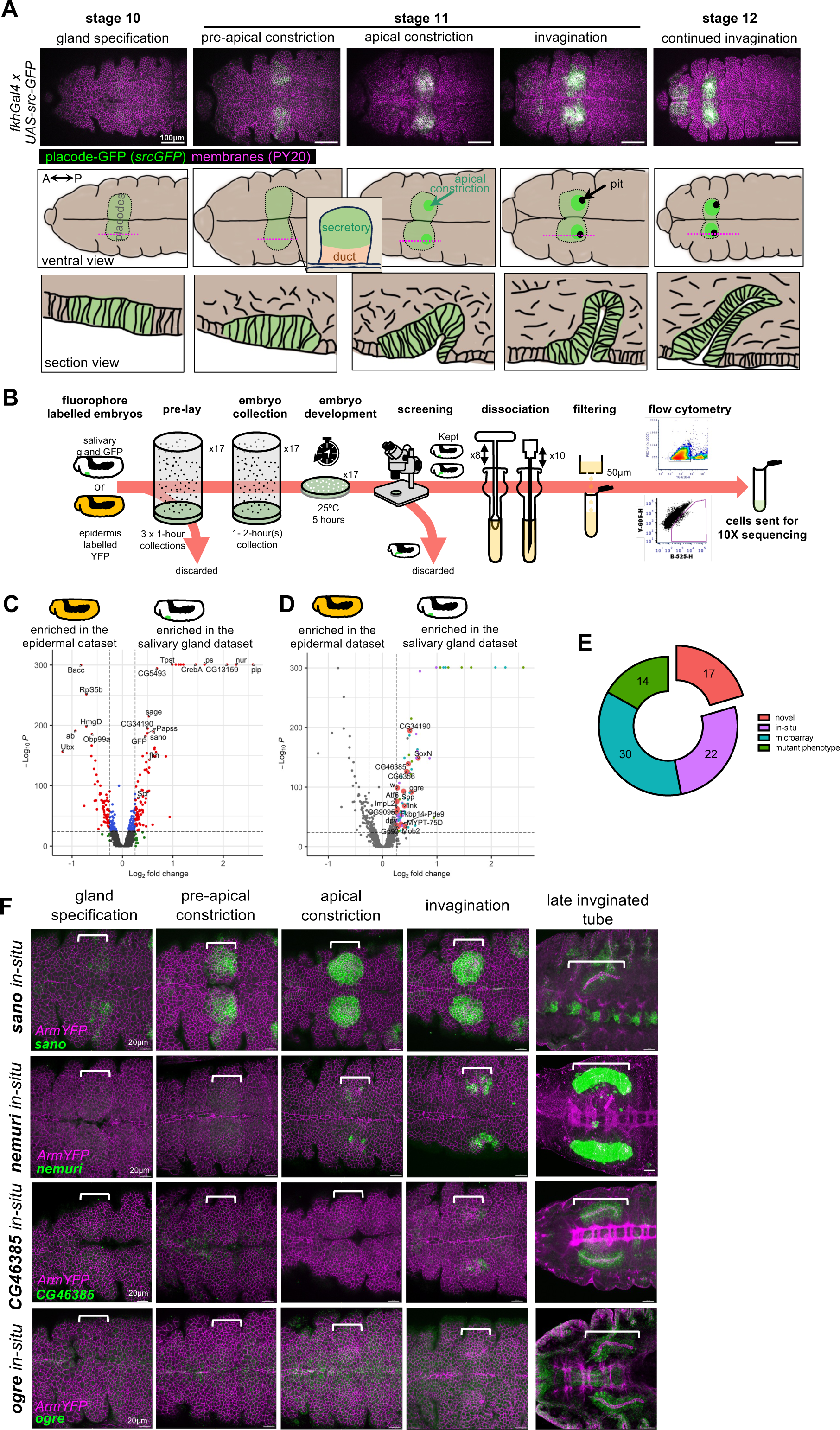
Generation of a single cell transcriptome dataset of salivary gland placodal and epidermal cells. **A** Immunofluorescence images of the anterior ventral half of *Drosophila* embryos at the indicated stages, highlighting the cells of the salivary gland placode, labelled with *fkhGal4 x UAS-srcGFP*. All cell outlines are labelled for phosphotyrosine to label adherens junctions (PY20, magenta) and srcGFP is in green. Scale bar is 100µm. Lower panels are matched schematics, showing the position of the salivary gland placode in pale green, the area of initial apical constriction in bright green and the forming invagination pit in black, as well as schematics of cross sections of the invaginating tube. **B** Schematic overview of the experimental pipeline for salivary gland placode (*fkhGal4 x UAS-srcGFP*) and epidermal (*armYFP*) cell isolation, FACS and 10X sequencing. **C-E** Pseudo-bulk differential expression analysis between salivary gland placodal (*fkhGal4 x UAS-srcGFP*) and epidermal (*armYFP*) cells, confirmatory genes highlighted in (**C**) and upregulated morphogenetic candidates highlighted in (**D**). **E** Genes specifically upregulated in salivary gland placodal cells were either known to show a phenotype in the glands when mutated (14), had been found to be expressed in the glands by whole-embryo microarray (30), had *in situ* images on public databases showing gland expression (22) or were completely novel with regards to expression or function in the salivary glands (17), colours are matched between **D** and **E**. **F** *In situs* by HCR for two novel identified genes upregulated in the salivary gland compared to the epidermis (*CG46385*, *ogre*) as well as for *nemuri* and *sano* that were also highly upregulated in our analysis in **D**. Cell outlines labelled by ArmYFP are in magenta and each respective *in situ* in green. Shown are representative images of early morphogenetic time points of the salivary gland placode as well as fully invaginated salivary glands. Scale bars are 50µm. Brackets indicate the position of the salivary gland placodes. See also Supplemental Figures S1 and S2.

In particular Fkh and Huckebein (Hkb) have emerged as transcription factors downstream of Scr that affect key aspects of the morphogenesis of the glands (Myat & Andrew, 2000a, b). In *fkh* mutants, no invagination or tube forms and Scr-positive cells remain on the surface of the embryo (Myat & Andrew, 2000a; Sanchez-Corrales *et al*., 2018). In *hkb* mutants by contrast cells invaginate, though in a central position and the forming glands have highly aberrant shapes (Myat & Andrew, 2000b; Sanchez-Corrales *et al*., 2021). Classical studies using embryo sections have suggested that the apical constriction and cell wedging at the forming invagination point is a key part of the internalisation of the tube, and that this is preceded by a lengthening of placodal cells and a repositioning of their nuclei towards the basal side (Myat & Andrew, 2000b). We have previously performed quantitative analyses of the live morphogenetic changes occurring in the early salivary gland placode, and identified two key additive cell behaviours, apical constriction or cell wedging near the forming invagination point as well as directional cell intercalation in cells at a distance to the pit. Furthermore, 4D investigation of cell behaviours also indicated further patterned changes with cells tilting and interleaving (Sanchez-Corrales *et al*., 2018). The apical constriction, we showed, is driven by a highly dynamic pool of apical-medial actomyosin (Booth *et al*, 2014). We identified that the apical constriction spreads out across the placode in a form of a standing wave, with intercalations feeding more cells towards the pit and cells switching from intercalation to apical constriction once they reach the vicinity of the pit. We could show that these changes are pre-patterned, at least in part, by dynamic transcriptional changes in the expression and protein distribution of the transcription factors Fkh and Hkb (Sanchez-Corrales *et al*., 2021).

Thus, not only tissue fate but also morphogenetic changes during tube budding are governed by transcriptional changes. Our previous analyses strongly suggest that the cell shape changes and behaviours that drive the tube budding morphogenesis are prepatterned transcriptionally in the tissue. We therefore decided to establish the transcriptional blueprint of the salivary gland placode just prior to and throughout the early stages of its morphogenesis using single cell genomic approaches, a blueprint required to initiate and drive the physical changes. Previous efforts to establish gene expression profiles across salivary gland tube morphogenesis included candidate *in situ* (Abrams & Andrew, 2005; Maruyama *et al*, 2011) as well as ChIPseq approaches (Fox *et al*, 2010; Johnson *et al*, 2020), whole embryo microarrays (Abrams & Andrew, 2005; Loganathan *et al*, 2016; Maruyama *et al*., 2011) and recently also whole embryo scRNAseq approaches (Calderon *et al*, 2022; Karaiskos *et al*, 2017; Peng *et al*, 2024; Seroka *et al*, 2022), though these latter approaches suffered from the fact that salivary gland placodal cells only constitute a very small fraction of all embryonic cells (and thus the datasets), or salivary gland placodal cells were not identified.

Here we generate and utilise a single cell RNA-sequencing dataset of isolated salivary gland placodal cells in comparison to isolated epidermal cells covering the early aspects of salivary gland tube morphogenesis. Pseudo-bulk differential expression analysis identifies a set of novel gland-specific factors, whereas the single cell analysis across pseudotime reveals complex regulatory patterns of expression. To our surprise, not only specific upregulation of factors either across the whole salivary gland primordium or within sections of it occurs, but in addition a number of factors become specifically downregulated and excluded in their expression from the placode, just at the start of morphogenesis. We identify that in certain cases this exclusion of expression is key to wild-type tube morphogenesis. In particular, we show that the ectopic continued expression of one of these factors, the <u>L</u>eucin-<u>R</u>ich <u>R</u>epeat receptor (LRR) Tollo/Toll-8, leads to aberrant morphogenesis, due to Toll-8 interference with endogenous systems, most likely the patterned expression and function of further LRRs, required for correct morphogenesis.

## Results

### Generation of salivary gland and epidermal single cell RNA sequencing datasets

In order to obtain a single cell RNA-sequencing dataset of salivary gland placodal cells covering the earliest stages from just after specification to early morphogenesis, as well as a matching epidermal cell dataset, we developed a new experimental pipeline: embryos of the genotypes *fkhGal4 x UAS-srcGFP* (for the salivary gland placodal cells; (Maybeck & Röper, 2009)) and *Armadillo/β-Catenin-YFP* (for the epidermal cells) were collected over a 1-2 hours period and aged for 5 hours before being subjected to further visual screening and selection in order to enrich for embryos of the desired stage (Fig.1 A, B). The *fkhGal4* driver used is based on a 1kb fragment of the *fkh* enhancer that drives expression very early in the salivary gland primordium, just downstream of specification (Zhou *et al*, 2001), and in combination with expression of a membrane-targeted form of GFP (Maybeck & Röper, 2009) highlights placodal cells early on (Fig. 1A and Suppl.Fig. S1A). The Armadillo-YFP fly stock contains a YFP-exon trap insertion into the *arm* locus, thereby labelling the endogenous protein (Lowe *et al*, 2014). Embryos were dissociated in a lose fitting Dounce homogeniser and filtered through a 50µm mesh to remove debris before being subjected to flow cytometry to sort GFP- or YFP-positive cells, and these were then subjected to 10X Chromium sequencing (Fig.1B, for details see Materials & Methods). Following quality control steps, including removal of doublets, dying and unspecified cells (Materials & Methods; Suppl.Fig. S2), a total of 3452 salivary gland placodal cells and 2527 epidermal cells were obtained and integrated into a single dataset for downstream analysis.

### Comparison of salivary gland placodal to epidermal gene expression at the onset of salivary gland tubulogenesis

We initially investigated differential expression of genes between the complete salivary gland placodal and epidermal datasets in a pseudo-bulk analysis to, firstly, benchmark and quality control both datasets and, secondly, identify novel upregulated candidates within the salivary gland placodal dataset (Fig. 1 C-E; Suppl.Table 1). Within the salivary gland dataset, we identified *GFP*, *fkh* and *Scr* as upregulated (*fkh* Log_2_Fold 0.535539503, p-value 8.50E-144, GFP Log_2_Fold 0.456845734, p-value 1.02E-182, *Scr* 0.311639192, p-value 2.98E-80), as would be expected from early placodal cells expressing a GFP label. Conversely, the epidermal dataset showed upregulated expression of *abdominal* (*ab*), a gene expressed only posteriorly to location of the salivary gland placode (Fig. 1C; Log2Fold -0.9333054, p-value 1.19E-191).

Analysis of the most upregulated genes within the salivary gland placodal in comparison to the epidermal dataset revealed 86 genes (Fig.1D). Of these, only 14 had previously been found to show salivary gland defects when mutants were analysed [*pipe* (Zhu *et al*, 2005), *pasilla* (Seshaiah *et al*, 2001), *CrebA* (Andrew *et al*, 1997), *Btk29/Tec29* (Chandrasekaran & Beckendorf, 2005), *PH4alphaSG2* (Abrams *et al*, 2006), *sage* (Chandrasekaran & Beckendorf, 2003; Fox *et al*, 2013), *myospheroid* (Bradley *et al*, 2003), *fkh* (Jürgens *et al*, 1984), eyegone (Isaac & Andrew, 1996; Kuo *et al*., 1996), *fog* (Lammel & Saumweber, 2000; Nikolaidou & Barrett, 2004), *Scr* (Mahaffey & Kaufman, 1987; Panzer *et al*, 1992), *KDEL-R* (Abrams *et al*, 2013), *crossveinless-c* (Kolesnikov & Beckendorf, 2007), *ribbon* (Bradley & Andrew, 2001)], 30 had been described to be expressed within the salivary gland by microarray analysis [*nemuri*, *CG13159*, *Hsc70-3*, *windbeutel*, *Papss*, *CG14756*, *PH4alphaSG1*, *SsRbeta*, *sallimus*, *TRAM*, *Sec61beta*, *twr*, *piopio*, *Spase12*, *PDI*, *CG7872*, *Surf4*, *Calr*, *CHOp24*, *par-1*, *p24-1*, *Trp1*, *Sec61gamma*, *nuf*, *RpS3A*, *Spase25*, *ERp60*, *Prosap*, *Fas3*, *fili*, (Fox *et al*., 2010; Loganathan *et al*., 2016; Maruyama *et al*., 2011)] and 22 could be identified to be expressed in the salivary glands or placode through publicly available *in situ* hybridisation databases [*Tpst*, *CG5493*, *sano*, *CG5885*, *Sec61alpha*, *Tapdelta*, *Gmap*, *l(1)G0320*, *nyo*, *NUCB1*, *Manf*, *bai*, *ergic53*, *CG32276*, *eca*, *GILT1*, *Spase22-23*, *Glut4EF*, *CG17271*, *bowl*, *CG9005*, *Tl;* (Hammonds *et al*, 2013; Lecuyer *et al*, 2007; Tomancak *et al*, 2002; Tomancak *et al*, 2007; Wilk *et al*, 2016). 17 genes we highly upregulated that had not been previously linked to salivary gland morphogenesis or function by any of the above means (*SoxN*, *ogre*, *CG34190*, *CG46385*, *CG6356*, *Mob2*, *link*, *Spp*, *MYPT-75D*, *Pde9*, *Fkbp14*, *Gp93*, *w*, *ImpL2*, *CG9095*, *Atf6*, *dpy*). To further validate these genes identified as salivary gland expressed, we performed *in situ* hybridisation using hybridisation chain reaction (HCR) for mRNAs of several genes that were either novel or where no spatio-temporal expression data existed, including *nemuri* that only began to be expressed in the salivary gland placode during onset of apical constriction, and *sano*, that began expression very early on in a region prefiguring the position of the forming invagination pit (Fig. 1F).

Thus, this differential expression analysis confirmed that our cell isolation and sequencing method was able to generate high quality datasets for further in depth analyses, and also revealed that we could identify novel expression of genes across a spread of early stages of salivary gland specification and morphogenesis.

### Generation of a salivary gland cell atlas by single cell RNAseq

We now focussed on the salivary gland dataset alone and used uniform manifold projection to identify clusters of cells with related expression profiles across this dataset. At a resolution of 0.17 the data split into 8 clusters (Fig. 2A; Suppl. Table 2). Analysis of top expressed genes predicted that these represented epidermal cells not yet specified to become salivary gland, salivary gland cells, anterior midgut cells, CNS cells, Enhancer of split [E(spl)]enriched cells, amnioserosa cells, muscle cells and hemocytes. The presence of cells other than salivary gland placode cells in this dataset was most likely due to the low level expression of the *fkhGal4* line also outside the salivary gland placode (see Fig. 1A), and also possible contamination due to mechanical dissociation applied to isolate cells. We aimed to confirm the identity of clusters and the expression or absence of top markers for each cluster in the salivary gland placode at stages when the morphogenesis had clearly commenced and performed HCR for these.

**Figure 2.**
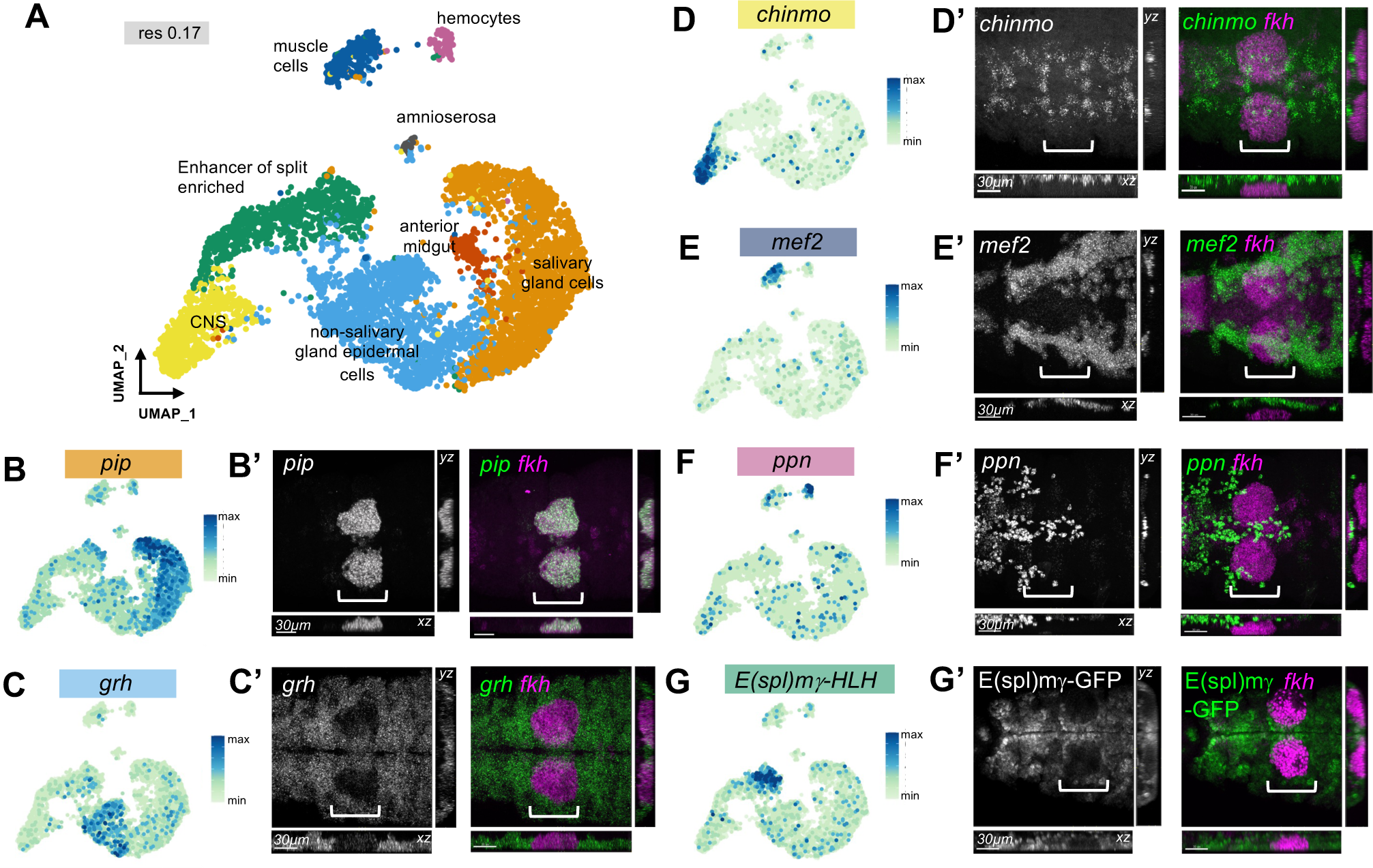
Single cell atlas of gene expression across early salivary gland morphogenesis. **A** Combined UMAP plot of all cells isolated in the *fkhGal4 x UAS-srcGFP* and *ArmYFP* samples, clustered at resolution 0.17. Salivary-gland specific and non-specific clusters are identified. **B-G’** Expression of specific marker genes for each cluster plotted onto the UMAP (**B-G**), with *in situ* hybridisation images by HCR for each gene shown in **B’-G’**. Marker genes are in green in merged channels, and also shown as individual channels. Images show face on views with matching xz and yz cross-sectional views, to distinguish labelling at different depth. White brackets indicate the position of the salivary gland placode. Scale bars are 30µm.

*pip* (*pipe*), a sulfotransferase of the Golgi is key to the later secretion function of the salivary glands and a known marker of these, and its transcript colocalised already early on with *fkh* mRNA, confirming this cluster as ‘salivary gland’ (Fig. 2B, B’). The top marker gene for the epidermal, pre-salivary gland cluster, *grh* (*grainyhead*), encoding a pioneer transcription factor (Hemphala *et al*, 2003; Jacobs *et al*, 2018; Narasimha *et al*, 2008) was expressed throughout the epidermis when analysed at stage 11 when apical constriction had commenced, not overlapping with *fkh* mRNA (Fig. 2C, C’). *chinmo*, encoding a transcription factor with a key timing role in the nervous system (Kao *et al*, 2012; Zhu *et al*, 2006), localised to groups of neuronal precursors, and although some of the expression looked to overlap *fkh* mRNA, 3D analysis revealed that *chinmo* expressing cells were in fact localised further interior than the salivary gland cells (Fig. 2D, D’). *mef2*, encoding a muscle transcription factor (Elgar *et al*, 2008), was expressed in muscle precursors at stage 11, localising further interior than *fkh* expressing placodal cells (Fig. 2 E, E’). *ppn* (*papilin*) encodes a components of the extracellular matrix (ECM)(Campbell *et al*, 1987) that is, as is most embryonic ECM, expressed by hemocytes during their embryonic migration and its expression therefore did not colocalise with *fkh* mRNA (Fig. 2 F, F’). *E(spl)* gene expression dominates one cluster (Couturier *et al*, 2019; Schrons *et al*, 1992), and analysis of a GFP trap in E(spl)mψ revealed protein expression across the epidermis but excluded from the salivary gland placode marked by *fkh* mRNA (Fig. 2 G, G’).

Thus, the single cell RNA-sequencing analysis of cells marked by and isolated based on GFP expression under *fkhGal4* control (*UAS-srcGFP fkhGal4*) was able to generate a cell atlas of the salivary gland placode as well as its precursor epidermis and nearby tissues at early stages of embryogenesis that provides a rich resource of expression data for these stages.

### Salivary gland specific clusters reveal temporally controlled expression of many factors potentially affecting morphogenesis

Increasing the resolution of clustering and homing in on salivary gland-related cells revealed a split into 4 clusters (Fig. 3A). Analysis of the highest expressed genes for each of these clusters allowed us to order them in a temporal progression representing aspects of salivary gland morphogenesis: ‘early salivary gland’, ‘specified secretory salivary gland cells’, ‘specified duct salivary gland cells’, ‘post specification salivary gland cells’ (Fig. 3 A, B; Suppl. Table 3). To understand better what set each cluster apart from the others, we looked at differential gene expression for each of these clusters (Fig. 3 B,C). Each cluster displayed a selective upregulation of genes, some of which had previously been linked to salivary gland morphogenesis or function, and others not implicated or known to be expressed in the placode or glands. We therefore performed *in situ* hybridisation to confirm the temporal changes in expression of marker genes for each cluster along the salivary gland morphogenesis trajectory, especially focussing on early stages of the process, split into ‘early’, ‘pre-apical constriction’, ‘apical constriction’ and ‘continued invagination’ (Fig. 3D). As previously described, *hth*, encoding a transcription factor working in conjunction with Scr and Exd in salivary gland placode specification (Henderson & Andrew, 2000), was expressed very early in the primordium and then appeared to be actively excluded from the placodal cells. Uncharacterized gene *CG45263*, a top marker gene within the specified duct cell cluster (Fig. 3B, C) was expressed in a spatial pattern that initiated close to the ventral midline with expansion into the duct cells of the salivary gland primordium during pre-apical constriction stages. Similar to the expression timing previously reported for components related to EGFR signalling within the duct cells of the salivary gland primordium (Zhou *et al*., 2001), expression of *CG45263* within the duct portion of the salivary glands continued even post-invagination (Fig. 3D). As previously described, mRNA for Gmap, a Golgi-microtubule associated protein, was expressed within the salivary gland placode beginning at early stage 11 (Friggi-Grelin *et al*, 2006). The *in situ* hybridisation covering early placodal development confirmed this observation and furthermore revealed that the expression originated in the cells first to invaginate at early stage 11, but that the onset of *Gmap* expression extending to all secretory cells was later than the onset of *fkh* expression (Fig. 3D). *calreticulin* (*calr*), the top marker gene of the ‘post specification’ cell cluster has not previously been implicated in a specific salivary gland development function. It showed a later onset of expression in the salivary gland placode than both *fkh* and *Gmap*, displaying a diffuse expression beginning during pre-apical constriction in all secretory cells with expression increasing as invagination continued.

**Figure 3.**
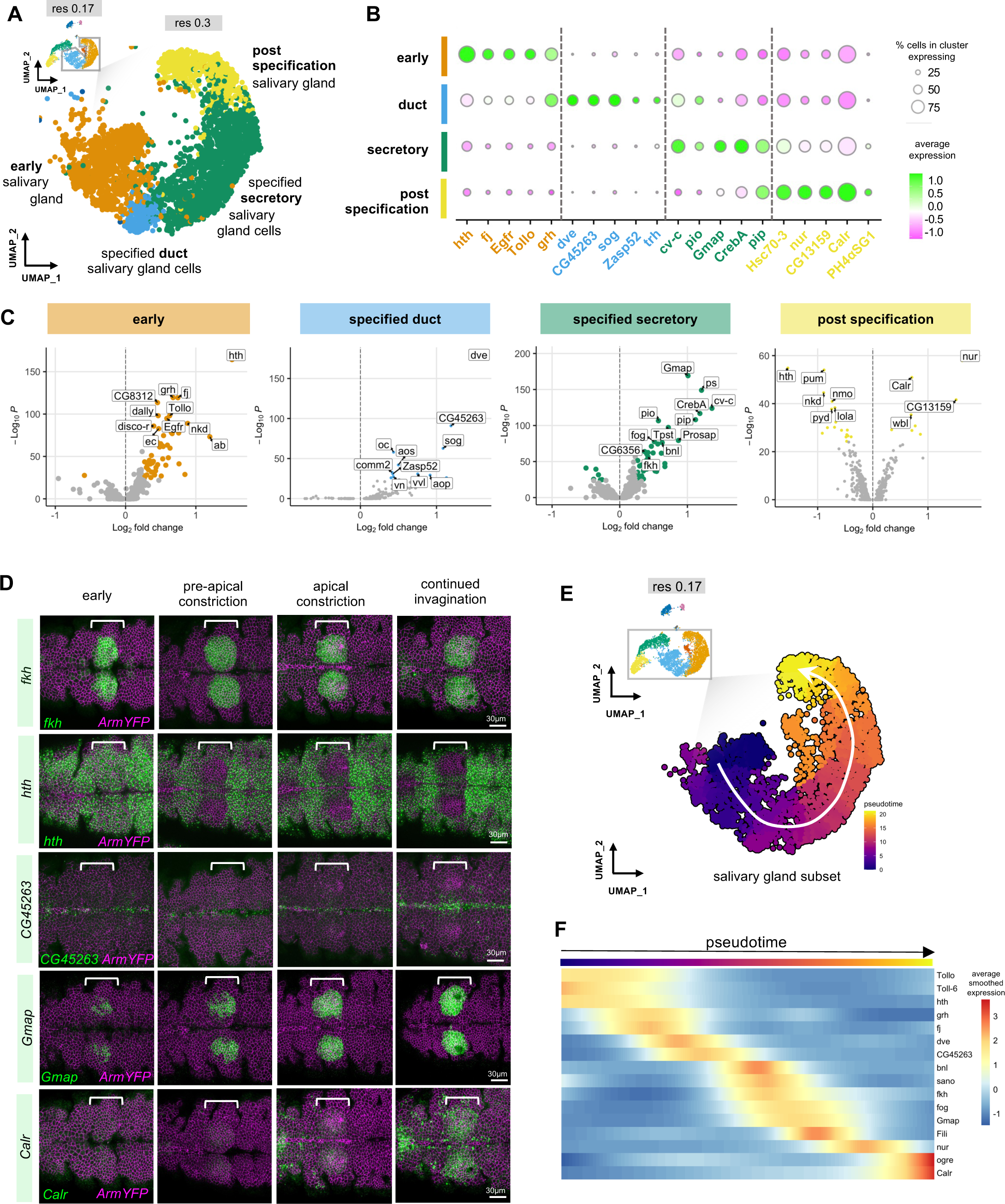
A single cell timeline of mRNA expression changes during salivary gland morphogenesis. **A** Higher resolution UMAP of the salivary gland lineage subsection (resolution 0.3), with clusters indicating a temporal progression along salivary gland morphogenesis labelled. **B** Expression analysis of the top five marker genes of each cluster identified in **A**: early salivary gland, specified duct cells, specified secretory cells and post specification salivary gland cells. Colour indicates level of expression and the size of the circle indicates the percent of cells in each cluster expressing the gene, vertical dotted lines denote cluster boundaries. **C** Volcano plots displaying upregulated marker genes in each of the clusters identified in **A**. **D** *In situ* hybridisation by HCR of one top marker gene per cluster identified in comparison to *fkh* expression: *hth* for the ‘early gland’ cluster, *CG45263* for the ‘specified duct cells’ cluster, *Gmap* for the ‘specified secretory cells’ cluster and *Calr* for the ‘post specification’ cluster. White brackets indicate the position of the salivary gland placodes, scale bars are 30µm, *in situ* for marker genes is in green, ArmYFP to label cell outlines is in magenta. **E** Pseudotime analysis based on the salivary gland portion of the lower resolution UMAP in Fig. 3A. **F** Based on the pseudotime trajectory a subset of genes was plotted along pseudotime to reveal differential temporal expression along the early morphogenetic timeline. See also Supplemental Figure S3.

Thus, our cluster analysis using increased resolution strongly suggested a temporally controlled pattern of gene expression along the morphogenetic trajectory. We now employed pseudotime analysis on the lower resolution cluster to analyse whether this approach would confirm our above analysis. Without specifying origin or endpoint clusters, unsupervised pseudotime analysis using Monocle3 identified a lineage originating from cells previously clustered in the ‘early salivary gland’ cluster moving towards the ‘post specification’ cell cluster (Fig. 3E). Focusing on this lineage, plotting the assigned pseudotime values of each cell on the previously generated UMAP indicated that cells analysed in this study are clustered along a temporal axis (Fig. 3E). This further validated the cluster assignment of cells in the salivary gland placode from early specification through to post-specification (Fig. 3A). Furthermore, the continuous change over time as indicated by the pseudotime analysis matched closely the biological reality of salivary gland tubulogenesis as a continuous process of invagination as opposed to discrete stages. Following the pseudotime trajectory also allowed us to identify and plot the differential expression of 207 genes, some of which are highlighted in Fig. 3F (Suppl. Table 4).

In summary, the expression analysis of placodal cells at the single cell level revealed an intriguing dynamicity of gene expression activation and cessation across the short time period of salivary gland morphogenesis, suggesting a tight temporal expression control of morphogenetic effectors.

### A cluster of genes with specific exclusion of expression in the salivary gland placode

In addition to the temporally controlled onset of expression of many factors within the salivary gland primordium, we also identified two groups of genes that showed a striking exclusion of expression during the stages spanning the tube morphogenesis. All of these genes, though, were strongly expressed in the primordium early on during or just after specification (Fig. 4).

**Figure 4.**
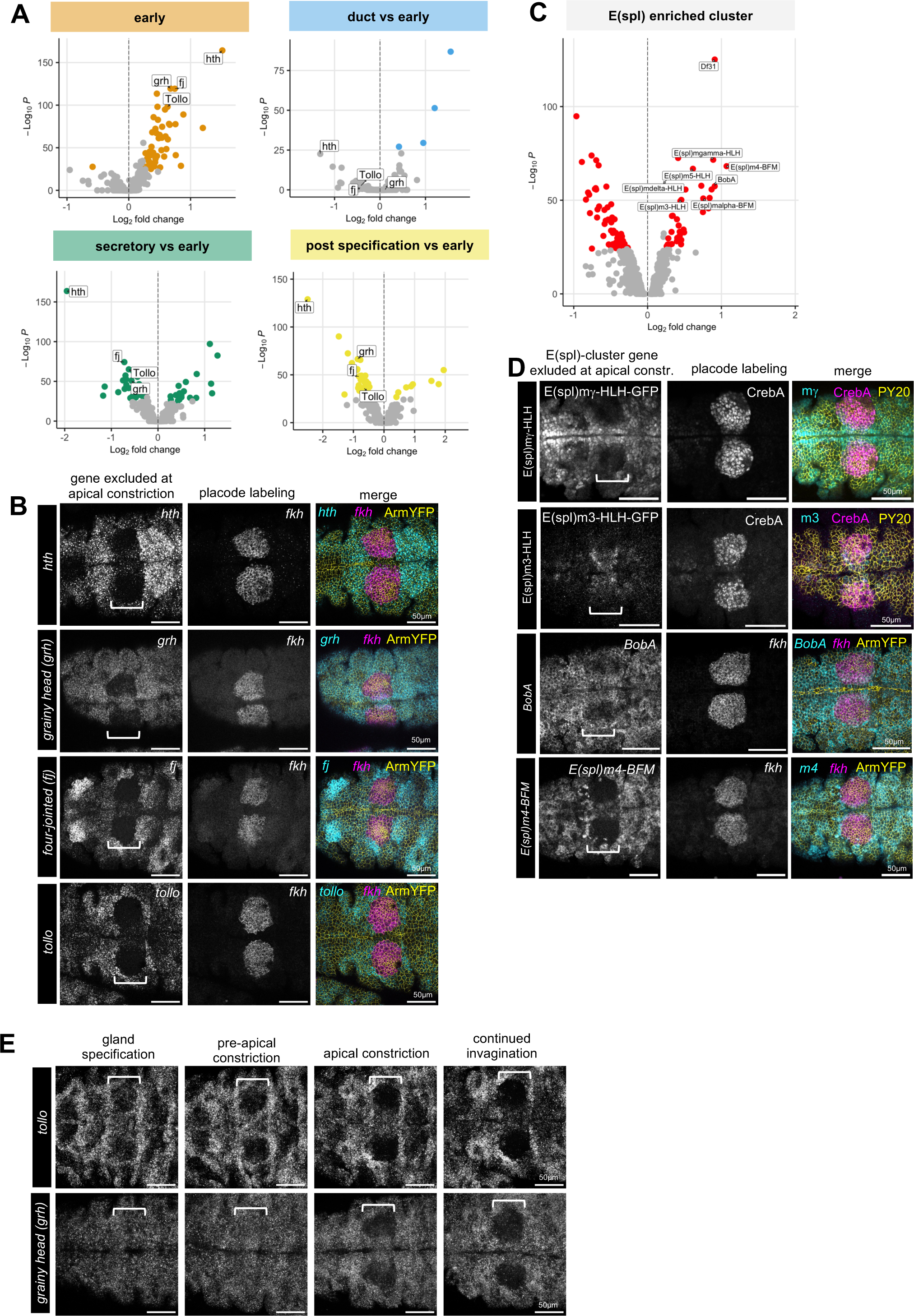
Placode-specific exclusion of expression of candidates. **A** Volcano plots showing candidates upregulated early during gland specification that become specifically downregulated in their expression once morphogenesis commences. **B** *In situ* hybridisation by HCR of salivary gland placodes for 4 downregulated genes at apical constriction stage (early stage 11, *hth*, *grh*, *fj* or *tollo* probes in cyan), in comparison to *fkh* in situ (magenta) and apical cell outlines (ArmYFP in yellow). **C** Volcano plot showing the E(Spl)-cluster that is also upregulated early during gland specification but then specifically downregulated and excluded during morphogenesis. Cut-off values are indicated by dashed lines. **D** Endogenously-tagged protein or *in situ* hybridisation by HCR of salivary gland placodes for 4 E(Spl) cluster genes (early stage 11, E(spl)mγ-HLH-GFP, E(spl)m4-BFM-GFP, *BobA* probe and *E(spl)m4-BFM* probe in cyan), in comparison to placode labeling via anti CrebA antibody or *fkh in situ* (magenta) and apical cell outlines (ArmYFP in yellow). **E** Timeline of *tollo* (top panels) and *grh* (lower panels) exclusion of expression in the salivary gland placode. Scale bars are 50 µm, white brackets indicate the position of the placodes.

The first group of genes comprised factors related to gland specification (*hth*) or, as previously described in other tissues, related to epithelial features (*grainy head* [*grh*], *four-jointed* [*fj*], *tollo*). Within the cell cluster defined above in the UMAP plot as containing cells of the early salivary gland (Fig. 3A; Suppl. Table 5), these four were strongly upregulated, but switched to being downregulated or excluded by the stage that morphogenesis commenced with apical constriction of cells to form the invagination pit (Fig. 4A). *In situ* hybridzation for *hth*, *grh*, *fj* and *tollo* revealed a mutually exclusive pattern of expression when compared to *in situ* labelling for *fkh* at the stage of apical constriction (Fig. 4B). In all cases the exclusion of expression was confined to the future secretory but not the future duct cells in the primordium (compare to *fkh* expression analysed in parallel).

The second group of genes all belonged to the Enhancer of Split [E(spl)] cluster, a group of transcriptional repressors involved in restricting neurogenic potential downstream of Notch in many tissues (Couturier *et al*., 2019; Schrons *et al*., 1992). *E(spl)m5-HLH*, *E(spl)m4-BFM*, *E(spl)mα-BFM*, *E(spl)m3-HLH*, *E(spl)mγ-HLH*, *E(spl)mδ-HLH* and *BobA* were all identified in this cluster (Fig. 4C). Also for this cluster, *in situ* hybridisation or use of GFP-reporter lines revealed a mutually exclusive pattern of expression when compared to *in situ* labelling for *fkh* or staining for CrebA, with the exclusion restricted to the future secretory cells (Fig. 4D).

For *tollo* and *grh*, two genes encoding factors previously implicated in either epithelial morphogenesis (*tollo*; (Lavalou *et al*, 2021; Pare *et al*, 2019; Pare *et al*, 2014)) or control of the epithelial phenotype and characteristics (*grh*; (Hemphala *et al*., 2003; Jacobs *et al*., 2018; Narasimha *et al*., 2008)), we analysed the spatio-temporal evolution of transcription for both over the time period of placode specification and early morphogenesis (Fig. 4E). At the gland specification stage, both *tollo* and *grh* were still expressed in parasegment 2 where the salivary gland placode will form, but both were clearly excluded from the secretory part of the placode once apical constriction commenced.

Thus in addition to the specific upregulation of factors within the salivary gland placode across specification and tube morphogenesis, it appears that there is a concomitant specific exclusion of expression of another set of factors, and this exclusion could represent another key part of the transcriptional programme driving tubulogenesis.

### Continued placodal expression of Tollo/Toll-8 reveals the requirement for its exclusion for correct salivary gland morphogenesis

The specific exclusion of *tollo* expression from the salivary gland placode coinciding with the onset of morphogenetic changes suggested that continued expression of *tollo* might interfere with these changes. We therefore decided to re-express Tollo specifically in the salivary gland placode under *fkhGal4* control. We used expression of both a full-length tagged version of Tollo (*UAS-TolloFL-GFP*) as well as expression of a tagged version lacking the intracellular cytoplasmic domain (*UAS-TolloΔcyto-GFP;* Suppl.Fig. S4). In control placodes (*fkhGal4*; Fig. 5B, bottom panels) with the start of apical constriction a narrow lumen tube started to invaginate and extend over time whilst cells internalised from the surface. By comparison when Tollo remained present across the placode (in *fkhGal4 x UAS-TolloFL-GFP* embryos), apical constriction appeared disorganised and spread to more cells. Already at early stages *fkhGal4 x UAS-TolloFL-GFP* embryos often showed the invagination of cells at multiple sites across the placode (Fig. 5B, arrows in cross sections), rather than the single wild-type invagination point. The invaginations then progressed to a too-wide and misshapen tube, and fully invaginated glands at stage 15 showed highly misshapen lumens and overall shape (Fig. 5B). Similar phenotypes were observed when TolloΔcyto was expressed (*fkhGal4 x UAS-TolloΔcyto-GFP; Suppl. Fig. S4*). In fact, compared to control placodes, *fkhGal4 x UAS-TolloFL-GFP* and *fkhGal4 x UAS-TolloΔcyto-GFP* placodes and invaginated tubes at stage 11 and 12 consistently showed multiple invagination points as well as too wide and branched lumens (Fig. 5C).

**Figure 5.**
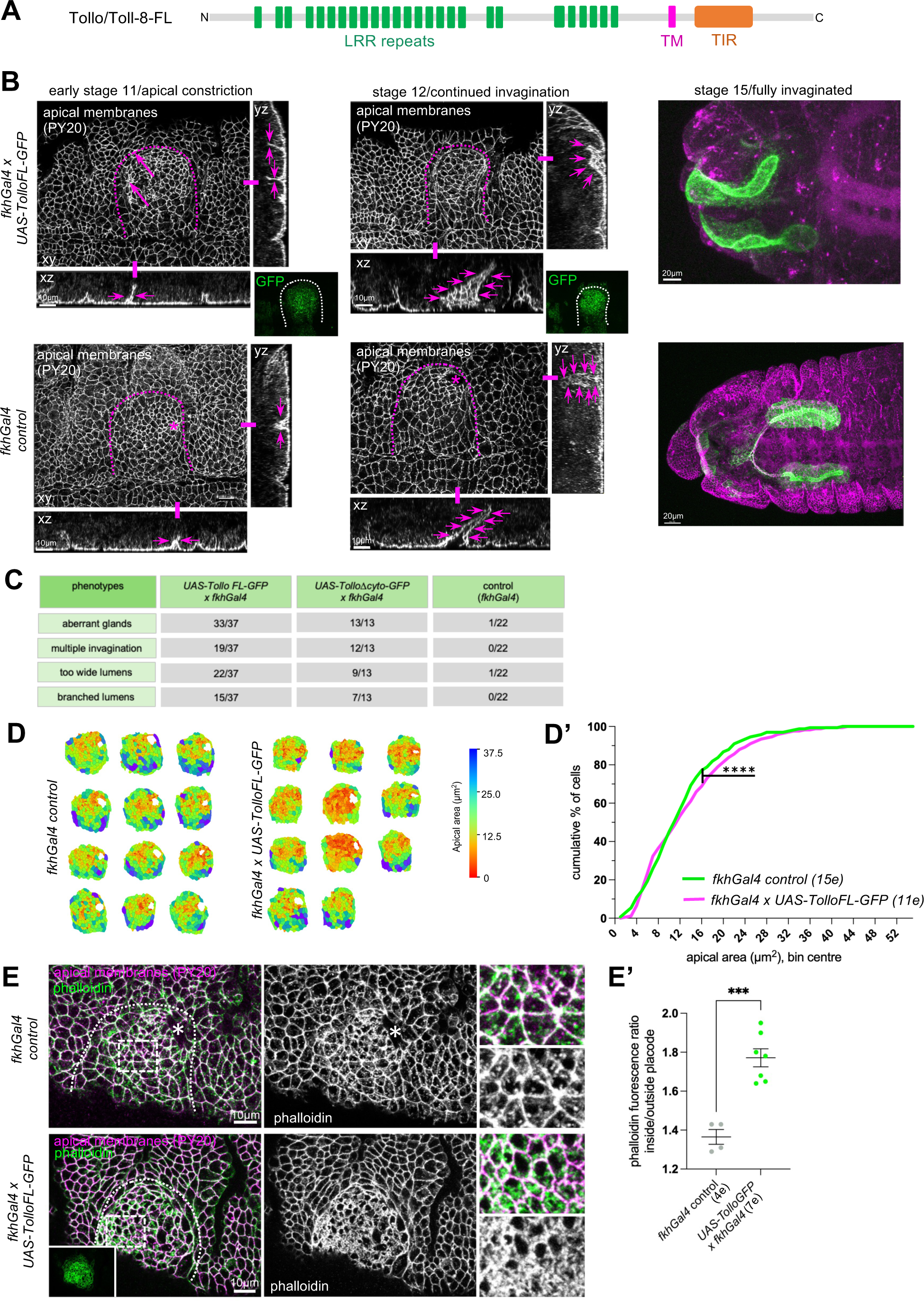
Continued expression of Tollo/Toll-8 disrupts salivary gland tubulogenesis. **A** Schematics of Tollo/Toll-8 full-length (FL) used for re-expression of Tollo/Toll-8 in the salivary gland placode using the UAS/Gal4 system. **B** In contrast to control (*fkhGal4 control*, bottom panels) placodes where apical constriction begins in the dorsal posterior corner and a narrow lumen single tube invaginates from stage 11 onwards, in embryos continuously expressing *UAS-TolloFL-GFP* under *fkhGal4* control (top panels) multiple initial invagination sites and lumens form and early invaginated tubes show too wide and aberrant lumens (magenta arrows in cross-section views). Fully invaginated glands at stage 15 show highly aberrant lumens. Apical membranes are labelled with an antibody against phosphotyrosine (PY20) labelling apical junctions. Dotted lines mark the boundary of the placode, asterisks the wild-type invagination point. Green panels show the expression domain of TolloFL-GFP. Scale bars are 10 µm or 20 µm as indicated. **C** Quantification of occurrence of aberrant glands, multiple invaginations, too wide lumens or branched lumens in either *fkhGal4 x UAS-TolloFL-GFP* or *fkhGal4 x UAS-TolloΔcyto-GFP* compared to *fkhGal4 control*. **D, D’** Quantification of apical area distribution of placodal cells in control (*fkhGal4 control*) and *fkhGal4 x UAS-TolloFL-GFP* placodes when invagination has commenced at stage 11. **D** Placode examples showing apical area. **D’** Quantification of the cumulative percentage of cells in different size-bins [Wilcoxon matched-pairs signed rank test, p<0.0001 (****)]. 12 placodes were segmented and analysed for control and 11 for *UAS-TolloFL-GFP* overexpression, the total number of cells traced was 1686 for control embryos and 1599 for *UAS-TolloFL-GFP* overexpression. **E, E’** Analysis of apical F-actin in placodes of *fkhGal4 control* in comparison to *fkhGal4 x UAS-TolloFL-GFP* labelled with phalloidin. Apical membranes are labelled for phosphotyrosine (PY20). Dotted lines mark the boundary of the placode, asterisks the wild-type invagination point. Green panel shows the expression domain of TolloFL-GFP. Scale bars are 10 µm. **E’** Quantification of apical phalloidin as a ratio of inside to outside placode in *fkhGal4 control* and *fkhGal4 x UAS-TolloFL-GFP* embryos. See also Supplemental Figure S4.

To address the aberrant apical constriction in a quantitative manner, we segmented the apical area of placodes in *fkhGal4* control and *fkhGal4 x UAS-TolloFL-GFP* embryos after apical constriction had commenced (Fig. 5 D, D’). This revealed an increase in apically constricted cells at stage 11 in embryos where Tollo continued to be expressed in the placode. We reasoned that, firstly, as we had previously shown that apical constriction depends on apical actomyosin in placodal cells (Booth *et al*., 2014), and, secondly, Toll/LRR proteins have been implicated in numerous contexts in the regulation of actomyosin accumulation at junctions (Pare *et al*., 2019; Pare *et al*., 2014)(Lavalou *et al*., 2021)(Tetley *et al*, 2016)(Peterson *et al*, 2023), the aberrant apical constriction could be due to changes in apical actomyosin. In *fkhGal4 x UAS-TolloFL-GFP* embryos, in comparison to *fkhGal4* control, apical F-actin was significantly enriched, in particular at junctions but also extending across large parts of the apical surface (Fig. 5 E, E’).

Thus, a continued presence of Tollo lead to significantly disrupted tubulogenesis of the salivary glands, strongly suggesting that the observed expression-exclusion of *tollo* is a key aspect of wild-type morphogenesis.

### Continued placodal presence of Tollo/Toll-8 interferes with endogenous LRR function in the placode

Why is the exclusion of *tollo* expression important for wild-type tube morphogenesis of the salivary glands? At earlier stages of morphogenesis during gastrulation three Tolls, Toll-2/18-wheeler, Toll-6 and Toll-8/Tollo, show an intricate alternate expression pattern across the epidermis that is key to germband extension movements (Pare *et al*., 2014). Furthermore, mutants in *toll-2/18-wheeler* (*18w*) have previously been reported to show defects in late salivary glands, a phenotype enhanced by further changes in components affecting the Rho-Rok-myosin activation pathway (Kolesnikov & Beckendorf, 2007). We therefore analysed the expression of *toll-2/18w* and *toll-6* in comparison to *toll-8/tollo* across early stages of salivary gland morphogenesis in the embryo (Fig. 6A,B). At early stage 10, just at the onset of salivary gland placode specification, all three genes were still expressed in a stripe pattern reminiscent of the earlier striped expression during gastrulation (Fig. 6A). At late stage 10/early stage 11, when the salivary gland placode was specified and morphogenesis was about to commence, *toll-2/18w* was expressed at the future invagination point in the placode, with expression radiating out form here, similar to what had been described previously (Kolesnikov & Beckendorf, 2007). By contrast, both *toll-6* and, as shown above, *tollo/toll-8* were now specifically excluded in their expression from the salivary gland placode (Fig. 6B). All three genes also showed more complex expression patterns in the epidermis surrounding the salivary gland placode at this stage.

**Figure 6.**
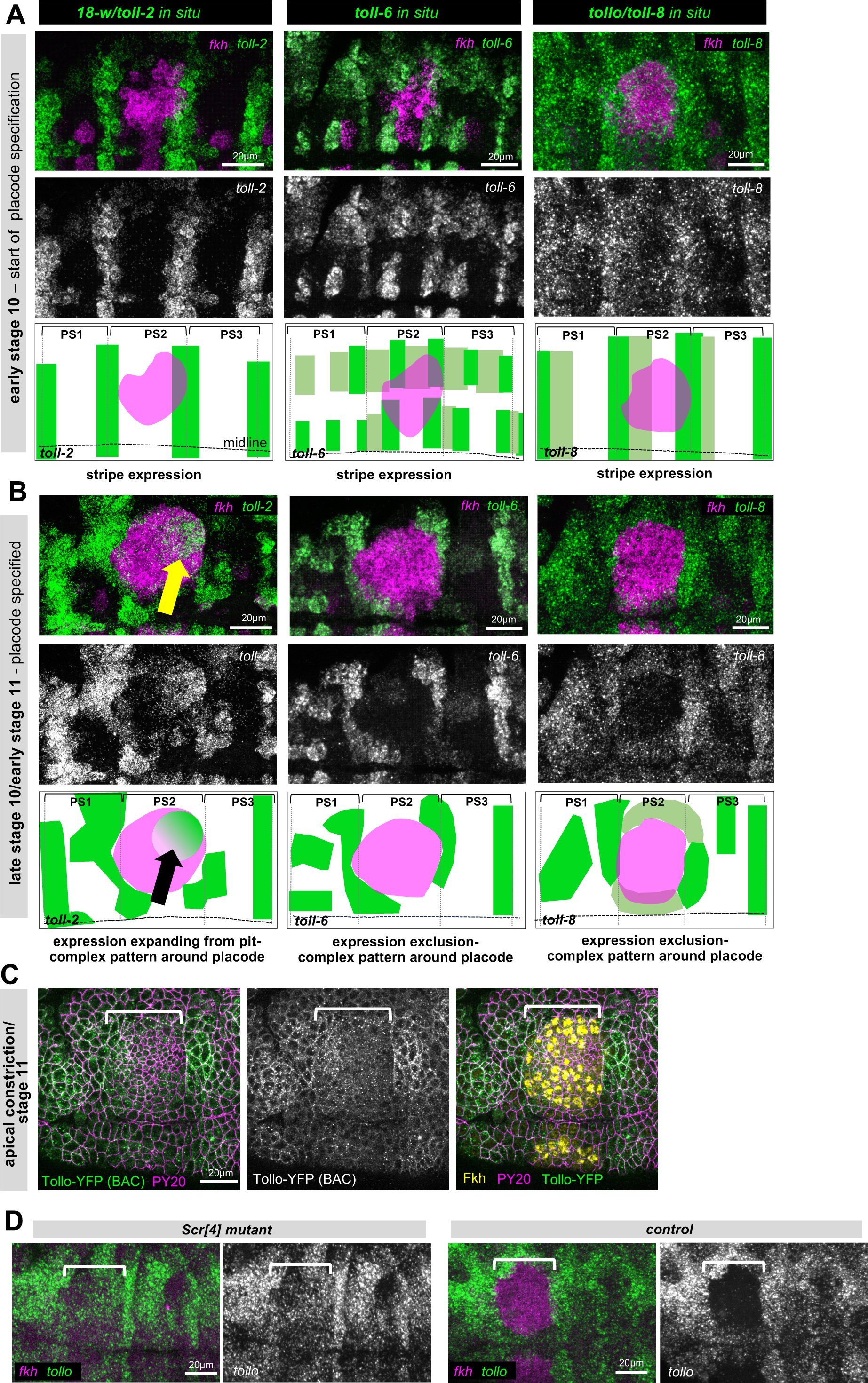
Continued expression of Tollo/Toll-8 disrupts an endogenous LRR code required for proper morphogenesis. **A** At early stage 10, prior to specification of the salivary gland primordium, *18-w/toll-2*, *toll-6* and *tollo/toll-8* are still expressed in complementary stripe patterns across the epidermis. Top row shows *in situ* hybridisations by HCR for each Toll in comparison to *fkh in situ*, lower panels show *toll-2*, *toll-6* and *toll-8* alone. The schematics show the stripe expression for each gene and *fkh* in comparison to the position of parasegmental boundaries. **B** At late stage 10/early stage 11 when apical constriction has commenced, *toll-2*, *toll-6* and *toll-8* show complex expression patterns with *toll-2* specifically expressed around the forming invagination point (yellow arrow) and *toll-6* and *toll-8* specifically excluded from the placode. The schematics show the stripe altered expression pattern for each gene and *fkh* in comparison to the position of parasegmental boundaries. **C** Tollo-YFP (*P{Tollo.SYFP2}*) protein expression in green at the apical constriction stage, apical cell outlines are marked using PY20 (anti-phospho-tyrosine) in magenta and placodal cells are marked by Fkh protein in yellow. **D** Comparison of *tollo/toll-8* expression analysed by *in situ* (HCR) in *Scr[4]* mutant embryos and control embryos at early stage 11. *tollo* is in green and *fkh* in magenta (note the expected absence of *fkh* in the *Scr[4]* mutant). Scale bars are 20µm, white brackets in **C** and **D** show the position of the salivary gland placode. See also Supplemental Figure S4.

We used BAC-mediated transgene expression of Tollo (Tollo-YFP) to analyse Tollo protein expression at early stages of salivary gland morphogenesis, with the caveat that this transgene in a wild-type background, though under control of endogenous elements, nonetheless leads to large fluorescent protein aggregates within the cells, likely due to the fact that cells now contain three copies of the *tollo* gene. We therefore focussed just on the apical surface of the placodal and epidermal cells at stage 11 when apical constriction had just begun. Tollo is usually localised to the apical junctional area (Lavalou *et al*., 2021). At early stage 11, Tollo-YFP appeared to be reduced in its levels within the salivary gland placode compared to the surrounding epidermis, matching its exclusion at the mRNA level (Fig. 6C).

Lastly, we wanted to test whether the specific exclusion of *tollo* expression from the salivary gland placode was part of the overall salivary gland morphogenesis programme initiated by the upstream homeotic transcription factor Scr or downstream Fkh. In *Scr[4]* mutant embryos at mid stage 11 *tollo/toll-8* expression was present across the salivary gland placode, when in the control in was already specifically excluded from it, indicating that Scr activity is responsible for the expression exclusion as part of the gland morphogenetic programme (Fig. 6D). By contrast, in *fkh[6]* embryos, *tollo/toll-8* expression occlusion still occurred (Suppl. Fig. 4C).

## Discussion

Complex forms during development arise from simple precursor structures, guided through detailed instructions that are laid down, at least initially, through a transcriptional blueprint and dynamic transcriptional changes. We know that for the tubulogenesis of the salivary glands in the fly embryo, transcriptional changes are key. Lack of the top-most transcription factors in the hierarchy, Scr, Hth and Exd, leads to complete lack of the glands and their primordia, but more revealing, overexpression of Scr leads to ectopic glands in anterior segments where Scr is not repressed, strongly suggestions that the whole morphogenetic cascade can be initiated by patterned expression of one factor alone in early embryogenesis (Andrew *et al*., 1994; Panzer *et al*., 1992). We also know that early morphogenetic macroscopic changes in the tissue are due to a delicate patterning of cell behaviours, including apical constriction and cell intercalation. Their patterning in the tissue primordium involves two key transcription factors (Sanchez-Corrales *et al*., 2018; Sanchez-Corrales *et al*., 2021). What was lacking was a more complete description of all transcriptional changes occurring in the placode primordium that could help guide our understanding of how other changes and behaviours are implemented that are key to the tubulogenesis and also later function of the tissue.

The single cell expression atlas of the salivary glands at early stages of morphogenesis presented here achieves this. Pseudo-bulk differential expression analysis identified many previously unknown factors whose role in the process will form the basis of future studies. The focus on the early stages of salivary gland tubulogenesis that this single cell atlas provides adds value to the study of this model process of tube budding, because other approaches at determining the transcriptional changing landscape of *Drosophila* embryos have limitations when it comes to salivary gland placodal cells. With a primordium of only about 200 cells in total within an embryo at stage 10-11 of about 48,000 cells, capturing enough salivary gland placodal cells in a whole embryo approach is always going to be challenging (Calderon *et al*., 2022; Karaiskos *et al*., 2017; Seroka *et al*., 2022)(Peng *et al*., 2024). Even if cells were captured, the stages analysed in whole embryo approaches usually did not span the time window we wanted to capture, i.e. the onset and early moprhogenesis. Hence, our single cell atlas provides a unique insight into this aspect of embryogenesis.

As we anticipated and aimed for, we identified many components that were expressed at specific early stages of gland morphogenesis, and that when analysed in *in situ* hybridisation showed gland specific expression either across the primordium (*pip, ogre, pasilla, fili*) or even in spatial patterns within the placode across time (*sano, fog, cv-c*) that we will explore for their significance in the future. Unexpectedly, we also identified a cluster of genes expressed prior to and during specification of the salivary gland placode that all then became specifically downregulated in and excluded from the placodal cells but not the surrounding epidermis (*hth*, *tollo*, *fj, grh*). This suggested that proteins encoded by these genes might in fact interfere with normal morphogenesis or specification of cells. For Hth, this exclusion of expression of its initial requirement for specification had been described (Henderson & Andrew, 2000). *hth* expression is both regulated by Scr but also together with ExD upstream of maintaining *Scr* expression in the placode early on. The trio of Scr/Hth/ExD is key for the expression of the next layer of transcription factors (Haberman *et al*., 2003), including Fkh and Hkb as two factors that are key for initiating part of the early morphogenetic changes (Myat & Andrew, 2000a, b; Sanchez-Corrales *et al*., 2021). Apart from the exclusion of *tollo* expression also being downstream of Scr but independent of Fkh, the further regulation of the specific exclusion of a whole set of genes is as yet unclear. As the exclusion was not dependent on Fkh, control must arise somewhere in between these two layers of regulation. Hairy as a transcriptional repressor has been shown to be important for salivary gland morphogenesis, thus far in regulating *hkb* and *crumbs* expression (Myat & Andrew, 2002). Disentangling the cascade of expression exclusion will broaden our knowledge of transcriptional control of morphogenetic events in the future.

The exclusion of expression of *tollo/toll-8* and *toll-6* is particularly intriguing as Toll/LRRs have been implicated in the regulation of junctional actomyosin accumulation by several studies, including in regulating germband extension in the early *Drosophila* embryonic epidermis (Lavalou *et al*., 2021; Pare *et al*., 2019; Pare *et al*., 2014). Furthermore, Toll-2/18w has been directly implicated in salivary gland morphogenesis, and mutants in *18w*, and especially zygotic double mutants in *18w* and *rhoGEF2* or *fog* show late gland phenotypes highly reminiscent of those we observed in the case of the continued expression of Tollo/Toll-8 in the placode (Kolesnikov & Beckendorf, 2007). Thus, the exclusion of *tollo/toll-8* expression could serve to prevent Tollo from interacting and interfering with 18w’s role, as LRR receptors can undergo homophilic or heterophilic interactions within this family (Özkan *et al*, 2013). Tollo/Toll-8, 18w/Toll-2 and Toll-6 have been shown to be able to interact heterophilically between neighbouring cells in S2 cell aggregation assays (Pare *et al*., 2014). The fact that continued placodal expression of the TolloΔcyto, i.e. Tollo lacking it cytoplasmic tail, resulted in the same phenotype as overexpression of the full-length version suggests that a possible interaction of the extracellular domain might titrate a required factor from its native interactions and this could be the reason for the morphogenetic problems. 18w/Toll-2 is not the only LRR expressed in the placode and involved in salivary gland tubulogenesis. We previously identified Capricious as an LRR protein whose overexpression results in a strong salivary gland tube defect (Maybeck & Röper, 2009). Using beta-galactosidase P-element traps we concluded that Capricious was endogenously expressed in tissues surrounding the salivary gland cells and Tartan, its usual interacting LRR (Mao *et al*, 2008), was expressed in placodal and gland cells itself. *tartan* and *capricious* double mutants show highly aberrant gland lumens (Maybeck & Röper, 2009). Thus, ectopic LRR expression in the placode could interfere with the endogenous expression and requirement of several LRRs.

In summary, our single-cell atlas of early salivary gland tube morphogenesis has already provided a rich source of identification of dynamically controlled expression or exclusion-of-expression of known or suspected morphogenetic effectors, and is likely to provide the basis for many further studies in the future.

## Materials and Methods

### Drosophila stocks and husbandry

The following fly stocks were used in this study:

*w;;fhkGal4 UAS-srcGFP* (Maybeck & Röper, 2009)*; Armadillo-YFP (PBac{681.P.FSVS-1}arm^CPTI001198^, w^1118^; ;)(Kyoto Stock Centre/DGGR); Tollo-YFP (BAC) (Pare et al., 2014); the following stocks were obtained from the Bloomington Drosophila Stock Centre: E(spl)mψ-HLH-GFP (#BL66401); E(spl)m3-HLH-GFP (#BL66402ß); UAS-TolloFL-GFP (#BL92990); UAS-Tolloτιcyto-GFP (#BL92991); Scr[4] (#BL942); fkh[6] (#BL545)*.

For expression in the placode, UAS stocks were combined with *fkhGal4* that is specifically expressed in the salivary placode and gland throughout development (Zhou *et al*., 2001).

### Embryo collection pipeline and Flow Cytometry

*Drosophila melanogaster* embryos expressing GFP in only the salivary gland (w*;;FkhGal4 UAS-srcGFP*) or expressing YFP in all epidermal cells (*PBac{681.P.FSVS-1}arm^CPTI001198^, w^1118^; ;*) were collected at 25°C in a humidity and CO_2_-controlled environment in 17 cages (973 cm^3^) per genotype. Three one-hour pre-lays were discarded prior to collection to reduce embryo retention in female flies and synchronise egg laying. Embryos were collected in one and two-hour time windows on apple juice agar plates with a small amount of yeast paste. Plates were removed from cages and incubated at 25°C for 5 hours 15 minutes. Embryos were washed into a basket and incubated in 50% bleach for 3 min for dechorionation and extensively washed. Embryos were removed from the basket and placed on cooled apple juice agar plates to slow developmental progression and visually screened using a fluorescence stereoscope with GFP filter. Embryos displaying an autofluorescent pattern indicating development beyond stage 12 were removed and only younger embryos up to this stage (10-12) were retained.

Mechanical dissociation of stage 10-12 embryos was performed using an adjusted method (Karaiskos *et al*., 2017). Approximately 5,000 embryos were placed in a 1 ml Dounce homogenizer containing 500 μl of ice-cold Schneider’s insect medium (Merk). Embryos were dissociated using 8 strokes of a loose pestle, followed by 10 gentle passes through a 16G 2-inch needle (BD microlance 3) into a 5 ml syringe. The final pass was filtered through a 50 μm filter (Sysmex) into a flow cytometry compatible tube. An additional 500 μl of ice-cold Schneider’s insect medium was added to the Dounce homogenizer and the process repeated to retrieve any additional cells to make a total of 1 ml of embryonic single cell suspension. To check single cell suspension was achieved, 20 µl of the suspension were mixed with an equal volume of trypan blue (Thermo), and placed on a CellDrop cell counter (DeNovix) to check live/dead ratios, and to visually assess that single cell suspension had been achieved and no large debris remained. Propidium Iodide (Thermo) was added to a final concentration 10 μg/ml and mixed into the suspension.

Single cells were sorted on a Sony Synergy system with gating for live cells, GFP signal and absence of autofluorescence signal. Live cells were identified by absence of propidium iodide signal, live cells were plotted on a secondary gate to isolate G/YFP+ cells from non-fluorescent cells and debris. This gate was determined by plotting signal from 488nm laser with a 525nm filter against signal from 488nm laser with a 510nm or 405nm filter. Prior to sorting, *yellow white* embryos were subject to an identical collection dissociation protocol and the boundary between non-GFP and G/YFP+ cells was determined by the maximum detected signal of *yellow white* embryos plotted on the same graph indicating non-GFP cells. G/YFP+ cells were collected in a 1.5 ml tube containing 37 μl of PBS and placed on ice before single cell sequencing. Three batches of cells were collected, two batches of *srcGFP* cells and one batch of *armYFP* cells. A total of 10,149 cells were collected for ArmYFP_1 and 10,000 for SrcGFP_1 and for SrcGFP_2 6,800 cells were submitted for sequencing to the Cancer Research UK sequencing facility for 10X library preparation and sequencing, according to the manufacturer’s protocols.

### Single cell RNA library preparation

Single cell suspensions were processed by Cancer Research UK, Cambridge, for 10X Chromium single cell sequencing. SrcGFP_1 and ArmYFP_1 were run on a singular lane of a NovaSeq flow cell at a 1:1 equimolar ratio with 10X v3.0 technology and sample preparation. SrcGFP_2 was run on a NovaSeq flow cell with two additional samples in an equimolar ratio of 3:1:3, the remaining two samples were excluded due to low cDNA quality. SrcGFP_2 was run on 10X v3.1 technology and sample preparation.

### RNAseq analysis

RAW FastQ files were obtained from the sequencing runs and a modified CellRanger 5.0.1 (10X Genomics) pipeline was applied to generate files for downstream analysis. Reads were aligned to a custom reference genome file generated using CellRanger mkref; this genome consisted of protein coding and antisense genes from the *Drosophila melanogaster* genome build 6.32 obtained from the FlyBase (www.flybase.org) FTP site and two additional custom gene transcripts (GFP and YFP). Firstly, a filtered GTF file was generated using CellRangers mkgtf package for custom reference genome building, the dmel6.32 GTF file from FlyBase, filtered for gene annotations selecting biotypes of protein coding and antisense genes. Following GTF file generation, GFP and YFP transcript sequences were obtained from f FlyBase, and added to the newly constructed GTF file. A reference genome file was then built using the custom GTF file and FASTA sequence file for the dmel6.32 genome build. Reads were aligned to the custom genome using CellRanger count. The original CellRanger outputs will be available on GEO.

### Seurat final object generation

All scripts run on CellRanger outputs and can be found in this paper’s GitHub repository [https://github.com/roeperlab/SalivaryGland_scRNAseq]. All sequencing analysis was carried out using R version 4.2.1 (R Core Team, 2022, https://www.R-project.org/), RStudio Build 576 (R Studio Team, 2020, http://www.rstudio.com/). CellRanger outputs were placed into Seurat objects, using the Seurat v4.1.3 package in R (Hao *et al*, 2021). Briefly, three samples were sequenced and used for subsequent analysis, two batches of salivary gland labelled cells and one batch of epidermal tagged cells. Prior to the merging of datasets the following limits were applied to all three datasets: cells containing more than 200 but less than 2,500 genes, cells containing less than 100,000 counts and cells containing less than 10% mitochondrial reads were retained in the dataset. Percentage mitochondrial reads were obtained by specifying mitochondrial genes as genes with the prefix “mt:”. An additional parameter of ribosomal gene percentage was obtained in a similar manner by specifying ribosomal gene reads to any gene beginning with “RpL” or “RpS”. Datasets were combined into a single Seurat object, and the remaining cell reads normalised using ScTransform with integrated anchoring methods, using 3000 genes to generate an anchor list (Stuart *et al*, 2019). Resolution of the combined dataset was decided via incrementally increasing resolution during FindAllClusters step and via plotting the splitting of clusters using Clustree v.0.5.0 (Zappia & Oshlack, 2018). For the generation of the total cell dataset, a further investigation into cells present in cluster 0, when including 44 dimensions at a resolution of 0.3, revealed no further clusters of relevance when increasing the resolution value. This cluster is likely a result of high RNA background within the dataset and cells assigned to this cluster were deemed low quality and removed from the dataset. The cells remaining from this pipeline are included in the Supplemental Information and a repeat analysis was carried out as above following the removal of low quality cells, the outcome of this analysis will be referred to as the complete cell dataset. The total dimensions used to generate the complete cell UMAP was 11 and plotted at a resolution of 0.17 and for the salivary gland development lineage was 0.3.

### Cluster identification

Initial cluster identification on the complete cell dataset was performed by running FindAllMarkers from the Seurat package, with literature reviews carried out for top markers of each cluster. In cases where cluster identity was not clearly assignable based on literature review or via tissue expression annotation in BDGP (Berkeley Drosophila Genome Project https://insitu.fruitfly.org/) or Fly-FISH (https://fly-fish.ccbr.utoronto.ca), in-situ hybridisation probes were obtained and the salivary gland region imaged across embryos at varying stages of salivary gland development in order to assign salivary or non-salivary gland identity.

For the more highly clustered salivary gland lineage, FindAllMarkers was used to generate a new marker gene lists for the newly identified clusters, a literature review was conducted and top markers from clusters were investigated in a similar manner. Top markers from this analysis and their respective Log_2_Fold change and adjusted p-values from this analysis were also used for DotPlot and VolcanoPlot generation. For additional comparisons between the early cluster and later clusters of the salivary gland lineage, Log_2_Fold change and adjusted p-values were obtained by running FindMarkers from the Seurat package, with the cluster of interest specified as ‘ident.1’ and the early cluster cells as ‘ident.2’. Genes of interest *hth, fj, Tollo, grh* and *E(spl)* cluster components were labelled on the resulting Volcano Plots. Volcano plots were generated using the R package EnhancedVolcano v.1.16.0 plotting the output of FindAllMarkers or FindMarkers gene marker tables.

### Pseudotime analysis

Pseudotime analysis was carried using a combination of Monocle3 (Cao *et al*, 2019), Slingshot (Street *et al*, 2018), and TradeSeq (Van den Berge *et al*, 2020) packages. The complete cell dataset was converted into a cell_dataset containing all metadata generated in Seurat including cell embeddings, reductions and cluster information to generate an object compatible with the Monocle package. The dataset was partitioned with a group label size of 3.5 to reflect the UMAP clustering (Levine *et al*, 2015). Using monocles learn_graph function, cells were assigned a pseudotime value and were ordered, root nodes were defined as the nodes covering the non-salivary gland epithelial cell population, prior to any predicted lineage splits. In order to reduce user bias final nodes were not specified. Cells were then plotted as a UMAP coloured by their assigned pseudotime value. For the generation of the pseudotime ordered heat map, cell embeddings from the complete cell dataset were used to generate a Slingshot object, a total of 6 lineages were identified with no starting or final cluster specified, of these 6 lineages, lineage 1 closely matched the predicted salivary gland lineage, all further analysis was carried out using this lineage. From lineage 1 a curve was generated using 150 points and a shrink value of 0.1 with 0 stretch. Using this curve pseudotime and cell weights were generated using Slingshot features slingPseudotime and slingCurveWeight, respectively. These values were then inputted into TradeSeqs NB-GAM model, by running fitGAM with 6 knots. Genes with high differential expression with respect to pseudtime were chosen for heatmap generation, their gene expression averaged and smoothed using predictSmooth from TradeSeq and the resulting data plotted in a heatmap using the pheatmap package (https://CRAN.R-project.org/package=pheatmap) with gene names and cells arranged via their pseudotime assigned value.

### Immunofluorescence analysis

Prior to immunoflourecene labeling embryos were fixed in a 4% formaldehyde solution and stored in 100% methanol at -20°C, or for embryos to be imaged with Rhodamine-phalloidin embryos were stored in 90% ethanol in water at -20°C. Embryos were rehydrated in PBT (PBS, 0.3% TritonX-100) followed by a five minute incubation in PBS-T (PBS, 0.3% TritonX-100, 0.5% bovine serum albumin) at room temperature. Embryos were blocked in PBS-T for a minimum of 1 hour at 4°C. Primary antibody solution was applied at varying concentrations (see reagent table) and incubated overnight at 4°C. The following morning primary antibody solution was removed and two washes in PBS-T were carried out at room temperature followed but 3 longer 20 minute washes before secondary antibody solution was applied and incubated for 1.5-3 hours. For immunoflourecene labeling containing Rhodamine-phalloidin, phalloidin was included in the secondary antibody solution. The secondary antibody solution was removed and a further two washes in PBS-T were carried out followed by three longer 20 minute washes before a final wash in PBS for five minutes. Embryos were mounted in Vectorshield (Vectorlabs H-1000) before being imaged. All immunofluorescence images were captured on an Olympus FluoView 1200 confocal microscope using a 40x oil objective.

### HCR analysis

Probes and hairpins for HCR in-situ hybridisation for genes of interest were obtained from Molecular Instruments, NM accession numbers were specified and in cases where genes had multiple isoforms regions of transcript shared amongst all transcripts were used to request probes. Embryos were collected and fixed in 4% Formaldehyde as detailed above, and stored in 100% methanol at -20°C before following a whole mount embryo HCR protocol (Choi *et al*, 2018). Batches of embryos were pooled and rehydrated in PBS + 0.1% Tween-20 (PBS-TW). Embryos were pre-hybridized at 37°C in hybridisation buffer followed by an overnight incubation at 37°C in primary probe solution (probe diluted to 0.8µM in hybridisation buffer). The following morning the probe solution was removed and embryos washed in probe wash buffer four times in 15 minute increments at 37°C followed by two short 5 minute washes in 5x SSCT buffer at room temperature. Secondary probes were chosen based on primary probe amplification region and the secondary probes’ emission signal. Care was taken to move any salivary gland probe markers to 647nm as to not overly saturate channels during imaging. Secondary probes were treated as individual hairpins (H1 and H2) initially and were separately heated to 97°C for 90 seconds before snap-cooling to room temperature. Embryos were amplified in room temperature amplification buffer for 10 minutes before combining H1 and H2 in amplification buffer to a final concentration of 0.8µM before being added to the embryos and incubated overnight in the dark at room temperature. The following morning secondary probes were removed and embryos washed in 5xSSCT for a 5 min wash followed by two 30 min washes and a final 5 min wash. All buffer was removed and embryos were mounted in VectaShield mounting medium containing DAPI (Vectorlabs H-1000) for immediate imaging. All *in situ* hybridisation images were captured as z-stacks on a Zeiss 710 Upright Confocal Scanning microscope with a 40x oil objective using full spectral imaging, and images were post-acquisition linear un-mixed. For linear unmixing using the Zeiss software individual spectra for each probe wavelength were obtained by carrying out in-situ hybridisation for highly expressing salivary gland genes (*fkh, CrebA*), one gene was chosen and one probe wavelength chosen. Regions of high signal were then specified and used to obtain spectral readings for the respective wavelength (488nm, 594nm or 647nm). *ArmYFP* embryos were scanned unstained to obtain the YFP spectrum and for the DAPI spectrum nuclei from *white* embryos mounted in VectaShield containing DAPI were used.

For early salivary gland development stages 10 and early stage 11 (pre-apical constriction, apical constriction) HCR was carried out on ArmYFP embryos in order to simultaneously image membrane labelling alongside mRNA expression, for these cases secondary probes at 488nm were not used.

### Quantification of apical area

For the analysis of apical cell area, images of fixed embryos of the genotypes w;;*fkh-gal4* embryos and w;;*fkh.gal4, UAS-TolloGFP* labelled with PY20 and phalloidin-Alexa633 were analysed. Images were categorised into apical constriction and continued invagination based on total cell numbers at the surface of the placode and surrounding morphological features. The first 4 optical sections (covering 4µm in depth) to display PY20 membrane signal were compiled into a maximum intensity projection and the PY20 membrane signal used for segmentation of the apical cell boundary. Segmentation was carried out with the Fiji plugin TissueMiner “*Detect bonds V3 watershed segmentation of cells*” (Etournay *et al*, 2016). The following parameters were used during segmentation: despeckling/ strong blur value of 3/ weak blur value of 0.3/ cells smaller than 5px excluded/ a merge basins criteria of 0.2/ kernel diameters in comparison to kuwahara pass and max pass of 5px and 3px, respectively. Automatic bond detection was hand-corrected to remove over-segmented cells and introduce missed bonds. Cells determined to be outside the placode were excluded. Cell data were exported from TissueMiner and reported cell areas were converted from px to µm^2^. Frequency distribution of cell areas were calculate and plotted using GraphPad Prism (version 9.5.1 for MacOS, GraphPad Software, Boston, Massachusetts USA). A histogram of the cell areas with bins of 2µm^2^ was generated and the values plotted as a cumulative percentage of cells.

### Quantification of apical F-actin

For the analysis of apical F-actin, images of fixed embryos of the genotypes w;;*fkhGal4*, w;;*fkhGal4; UAS-TolloGFP* were stained with PY20 to label apical cell outlines and Rhodamine-phalloidin to label F-actin. To assess the difference in phalloidin signal (F-actin) across the apical placodal area, for each analysed placode/embryo, using projections of the apical-most 4 confocal sections (each of 1µm thickness), three 10µm x 10µm areas both inside and outside the placode were quantified for fluorescence intensity per area, averaged and expressed as an inside/outside-ratio that was plotted.

### Statistical analysis

For comparison of cumulative distributions in the analysis of apical area size a Wilcoxon matched-pairs signed rank test was used, the relevant p value is indicated in the figure legend. For analysis of apical F-actin enrichment compared between different genotypes, differences in ratio values for fluorescence intensity inside/outside placode were tested for significance using un-paired Student’s t-test, mean +/- SEM are plotted and p-value is indicated in the figure legend.

## Data availability

The datasets and computer code produced in this study are available in the following databases:

- RNA-Seq data: Gene Expression Omnibus (upload pending)
- scRNAseq analysis computer scripts: GitHub (https://github.com/roeperlab/SalivaryGland_scRNAseq)

## Acknowledgments

The authors would like to thank the following people; for reagents and fly stocks: Debbie Andrew, Jen Zallen, Thomas Lecuit, Sarah Bray. Stocks obtained from the Bloomington Drosophila Stock Center (NIH P40OD018537) were used in this study and we thank them for their efforts.

This work was supported by the Medical Research Council, as part of United Kingdom Research and Innovation (also known as UK Research and Innovation) [MRC file reference number MC_UP_1201/11]. For the purpose of open access, the MRC Laboratory of Molecular Biology has applied a CC BY public copyright licence to any Author Accepted Manuscript version arising’.

## Author contribution

Conceptualisation, K.R. and A.M..; Methodology, K.R., A.M.; Investigation, K.R., A.M; Writing-Original Draft, K.R., A.M.; Funding Acquisition, K.R.

## Supplementary

**Supplemental Figure S1, related to Figure 1.**
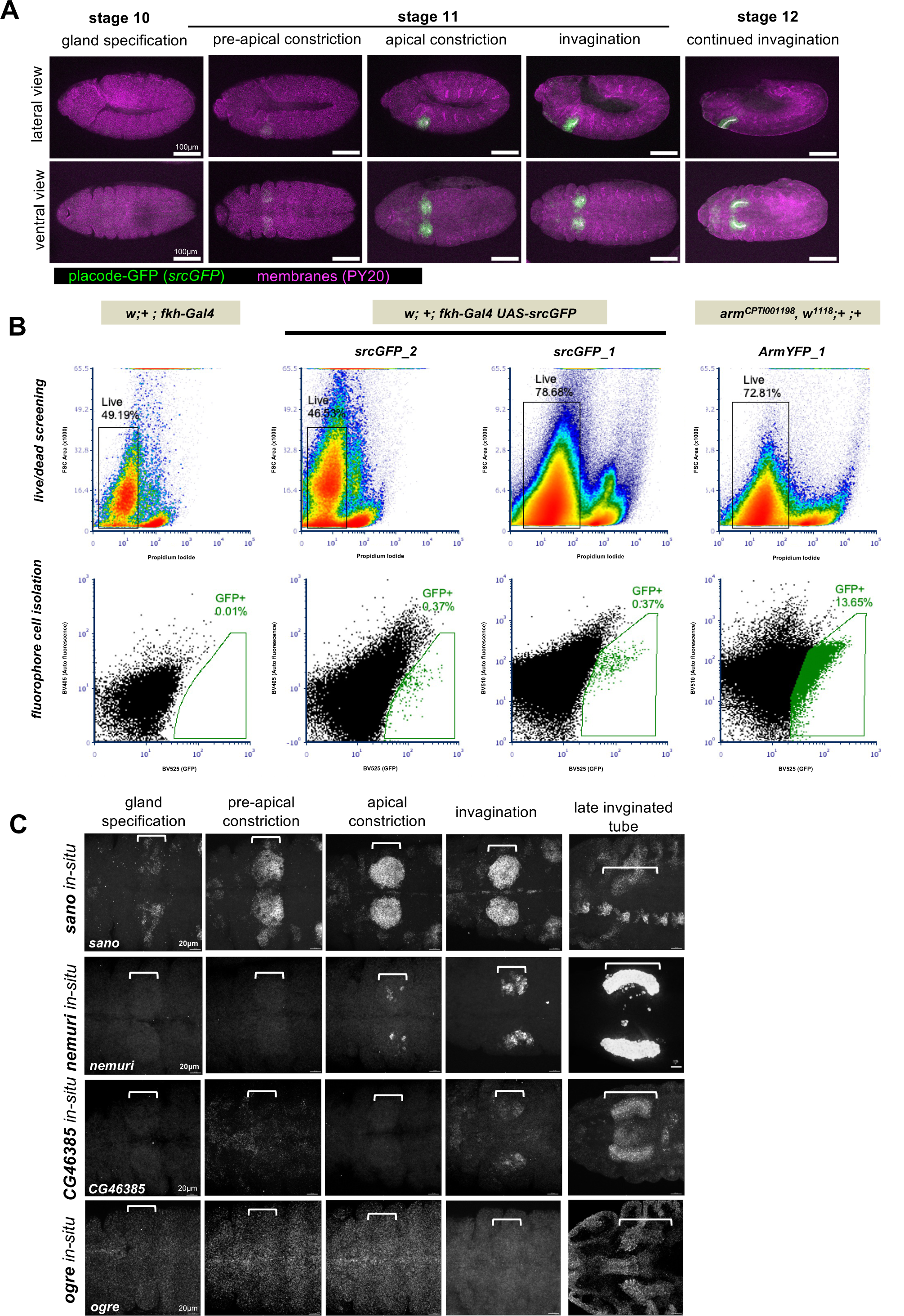
Generation of a single cell transcriptome dataset of salivary gland placodal and epidermal cells. **A** Whole embryo lateral (top row) and ventral views of *Drosophila* embryos at the indicated stages, highlighting the cells of the salivary gland placode as labelled with *fkhGal4 x UAS-srcGFP*, illustrated are early stages of morphogenesis. All cell outlines are labelled for phospho-tyrosine to label adherens junctions (PY20, magenta) and srcGFP is in green. Scale bars are 100µm. **B** FACS plots for non-fluorescent control (*w;+;fkhGal4*), two srcGFP embryo batches (srcGFP_1 and srcGFP_2; *w;fkhGal4 UAS-srcGFP*) and one ArmYFP embryo batch (*arm[CPTI001198], w[118];+;+*) used for single cell RNAsequencing. The top row shows the sorting for live vs dead cells, the bottom row shows the gate for sorting of GFP/YFP-positive cells and the percentage of total cells sorted they comprised. **C** Single channel of in situ hybridisation by HCR probes for the indicated genes. Matching two channel images are shown in Figure 1F.

**Supplemental Figure S2, related to Figure 1.**
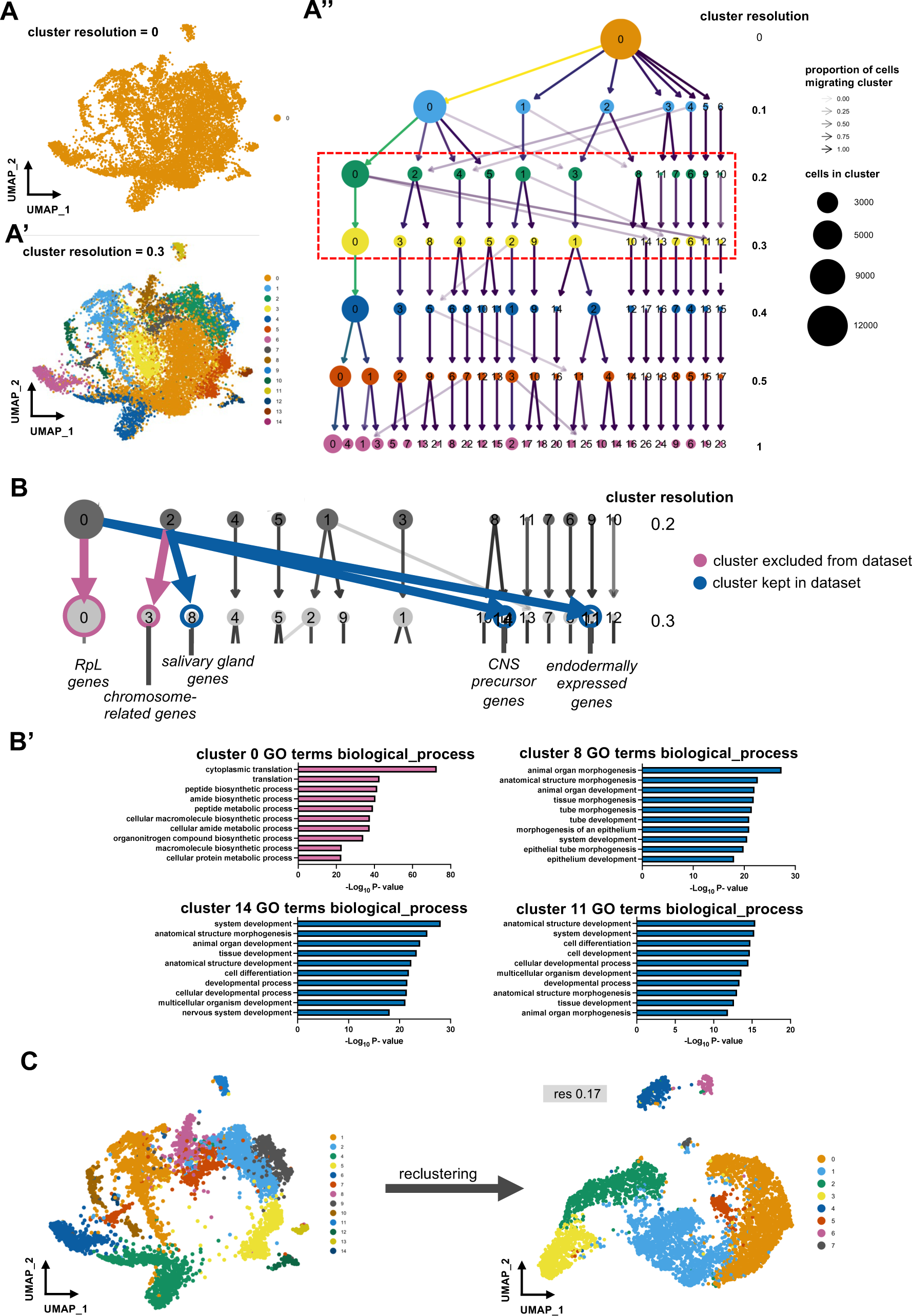
Generation of a single cell transcriptome dataset of salivary gland placodal and epidermal cells. **A-A’’** UMAP of single cell RNA sequencing at cluster resolution 0 (**A**) and 0.3 (**A’**). **A’’** Illustrates the emergence and linkage of clusters with increasing resolution generated using the Clustree package in R. Red dotted outline shows clusters assigned at resolutions of 0.2 and 0.3 where further investigations into cluster makers occurred. **B** General consensus of markers represented in the clusters emerging at resolution 0.3, purple arrows represent the emergency of cell clusters with markers of low-quality cells and blue arrows represent the emergence of clusters with biologically relevant cell types. **B’** Biological process Gene Ontology terms for gene lists generated from resolution 0.3 clusters featured in **B**, ranked by the -Log_10_P-value provided by FlyMine curated lists for each ontology term. **C** UMAP generated following the reclustering of the original dataset following the exclusion of low-quality cell types identified in **B**.

**Supplemental Figure S3, related to Figure 3.**
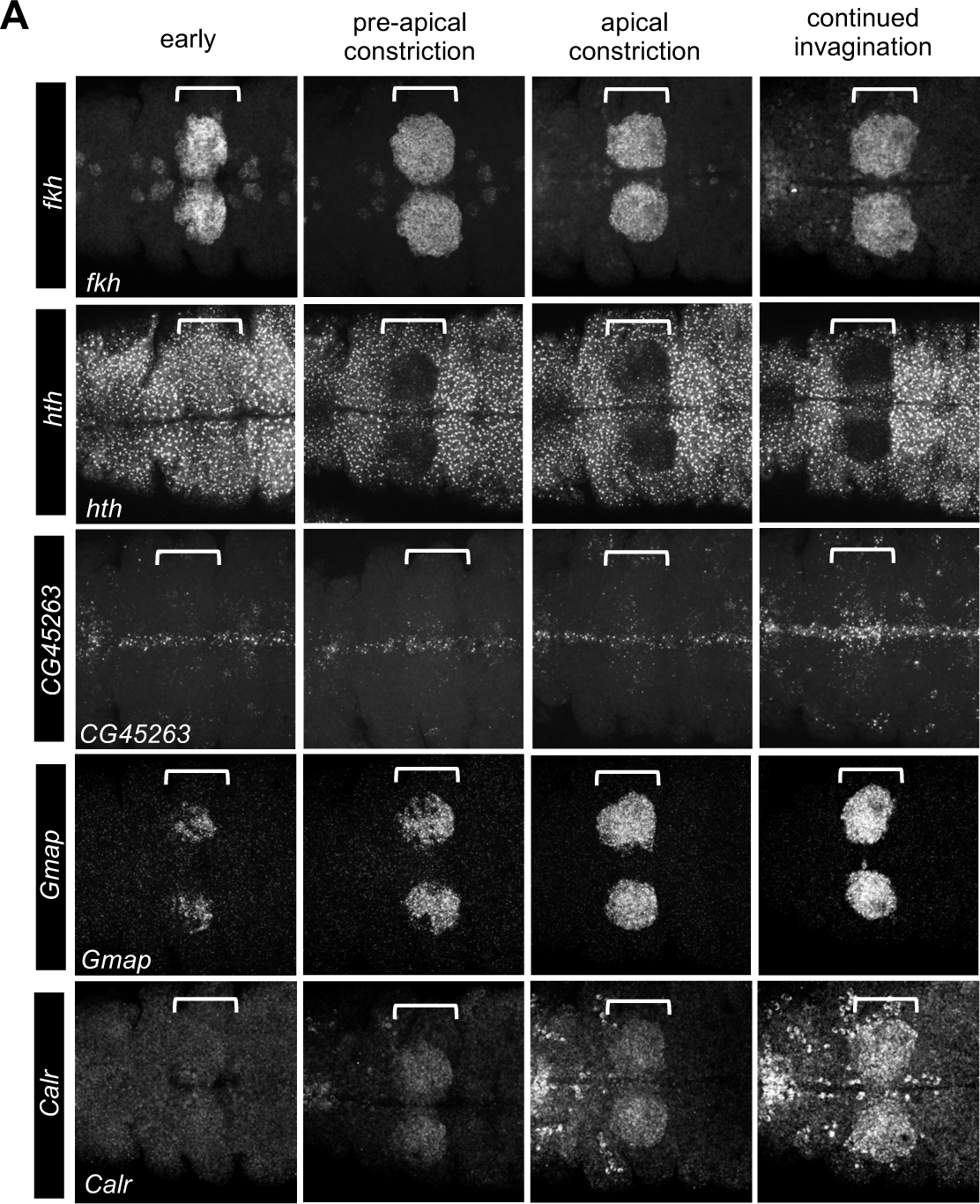
A single cell timeline of mRNA expression changes during salivary gland morphogenesis. **A** *In situ* hybridisation by HCR of one top marker gene per cluster identified in comparison to *fkh* expression as shown in Fig. 3D, single channels are shown here: *hth* for the ‘early gland’ cluster, *CG45263* for the ‘specified duct cells’ cluster, *Gmap* for the ‘specified secretory cells’ cluster and *Calr* for the ‘post specification’ cluster. Single in situ channels matching the panels in Figure 3D are shown. White brackets indicate the position of the salivary gland placodes, scale bars are 30µm.

**Supplemental Figure S4, related to Figure 5 and 6.**
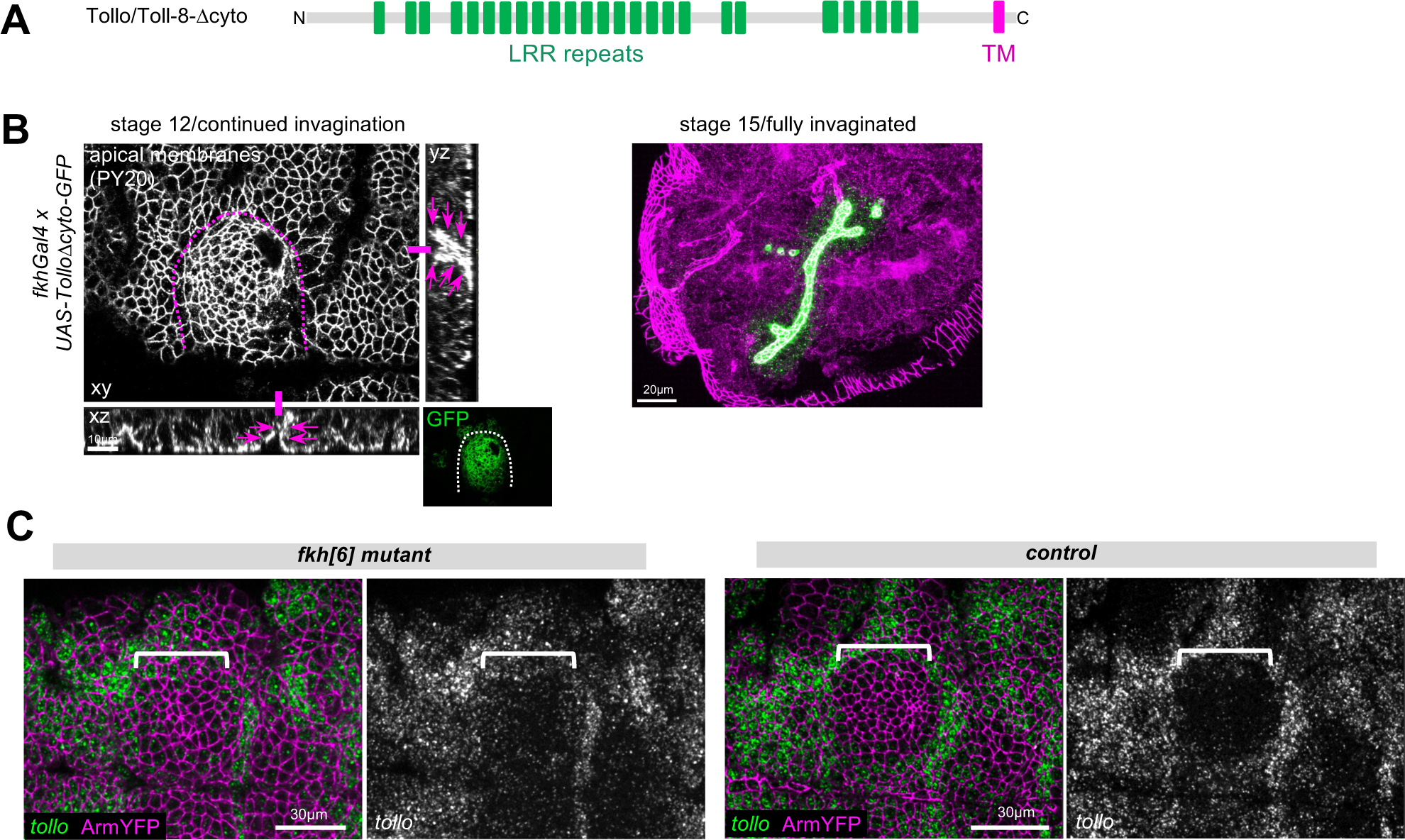
Continued expression of Tollo/Toll-8 disrupts salivary gland tubulogenesis. **A** Schematics of Tollo/Toll-8 lacking the intracellular cytoplasmic domain (Δcyto) used for re-expression of in the salivary gland placode using the UAS/Gal4 system. **B** In contrast to control placodes (Figure 5B) where apical constriction begins in the dorsal posterior corner and a narrow lumen single tube invaginates from stage 11 onwards in embryos continuously expressing *UAS-TolloΔcyto-GFP* under *fkhGal4* control multiple initial invagination sites and lumens form and early invaginated tubes show too wide lumens (magenta arrows in cross-section views). Fully invaginated glands at stage 15 show highly aberrant lumens. Apical membrane are labelled with an antibody against phosphotyrosine (PY20) labelling apical junctions. Dotted lines mark the boundary of the placode, asterisks the wild-type invagination point. Green panel show the expression domain of TolloFL-GFP. **C** Comparison of *tollo/toll-8* expression analysed by *in situ* (HCR) in *fkh[6]* mutant embryos and control embryos at stage 11. *tollo* is in green and ArmYFP in magenta. Scale bars are 30µm, white brackets in **C** show the position of the salivary gland placode.

**Supplemental Table 1, related to Figure 1C.** List of marker genes identified for cells of salivary gland placodal origin and epidermal origin and accompanying information. Marker genes generated by comparing cells originating from *fkhGal4 x UAS-SrcGFP* embryos sorted for GFP signal (salivary gland placode) and cells originating from *ArmYFP* embryos sorted for YFP signal (epidermis/epithelium) scRNAseq data, as plotted in Figure 1C. Limits applied to dataset for literature review categorisation were p-value adjusted ≤ 10E-25 and Log2Fold change ≥ 0.25 or ≤ -0.25. Genes categorised by highest piece or relevant information in the following order: 1) mutant phenotype in the salivary gland (MP), 2) microarray data showing expression or altered expression in the salivary gland in mutant phenotypes (MA), 3) *in situ* database images displaying expression in the salivary gland (I) or 4) no available information on expression in the salivary gland (New).

**Supplemental Table 2, related to Figure 2A.** Marker genes for the cell type clusters of combined salivary gland and epithelial cells. Marker genes for each individual cluster of the combined dataset at a resolution of 0.17 (see Figure 2A), generated using scRNAseq data. Limits applied to dataset: p-value adjusted ≤ 10E-25 and Log2Fold change ≥ 0.25 or ≤ -0.25. Each cell type cluster is displayed on an individual sheet, additionally data from the E(spl)-enriched sheet is plotted in Figure 4C.

**Supplemental Table 3, related to Figure 3A.** Marker genes defining clusters within the salivary gland temporal lineage. Marker genes for each individual subcluster of the salivary gland lineage at an increased resolution of 0.3 (see Figure 3A), generated using scRNAseq data. Limits applied to dataset: p-value adjusted ≤ 10E-25 and Log2Fold change ≥ 0.25 or ≤ -0.25. Each cell type cluster is displayed on an individual sheet.

**Supplemental Table 4, related to Figure 3F.** Differentially expressed genes across pseudotime in the salivary gland lineage. List of genes which are differentially expressed according to their wald statistic value (waldstat_1), p-value and mean log2fold change across the salivary gland lineage (see Figure 3F) generated from scRNAseq data. Limit: p-value < 0.05.

**Supplemental Table 5, related to Figure 4A.** Marker genes for clusters within the salivary gland temporal lineage when compared to the earliest cluster. Marker genes for each individual subcluster of the salivary gland lineage at a resolution of 0.3 when compared to the earliest cluster of cells (see Figure 4C), generated using scRNAseq data. Limits applied to dataset: p-value adjusted ≤ 10E-25 and Log2Fold change ≥ 0.25 or ≤ -0.25. Each cell type comparison is split into an individual sheet.

## References

1. Abrams EW, Andrew DJ (2005) CrebA regulates secretory activity in the Drosophila salivary gland and epidermis. Development 132: 2743–2758

2. Abrams EW, Cheng YL, Andrew DJ (2013) Drosophila KDEL receptor function in the embryonic salivary gland and epidermis. PLoS One 8: e77618

3. Abrams EW, Mihoulides WK, Andrew DJ (2006) Fork head and Sage maintain a uniform and patent salivary gland lumen through regulation of two downstream target genes, PH4alphaSG1 and PH4alphaSG2. Development 133: 3517–3527

4. Abrams EW, Vining MS, Andrew DJ (2003) Constructing an organ: the Drosophila salivary gland as a model for tube formation. Trends Cell Biol 13: 247–254

5. Andrew DJ, Baig A, Bhanot P, Smolik SM, Henderson KD (1997) The Drosophila dCREB-A gene is required for dorsal/ventral patterning of the larval cuticle. Development 124: 181–193

6. Andrew DJ, Horner MA, Petitt MG, Smolik SM, Scott MP (1994) Setting limits on homeotic gene function: restraint of Sex combs reduced activity by teashirt and other homeotic genes. Embo J 13: 1132–1144

7. Booth AJ, Blanchard GB, Adams RJ, Röper K (2014) A dynamic microtubule cytoskeleton directs medial actomyosin function during tube formation. Dev Cell 29: 562–576

8. Bradley PL, Andrew DJ (2001) ribbon encodes a novel BTB/POZ protein required for directed cell migration in Drosophila melanogaster. Development 128: 3001–3015

9. Bradley PL, Myat MM, Comeaux CA, Andrew DJ (2003) Posterior migration of the salivary gland requires an intact visceral mesoderm and integrin function. Dev Biol 257: 249–262

10. Calderon D, Blecher-Gonen R, Huang X, Secchia S, Kentro J, Daza RM, Martin B, Dulja A, Schaub C, Trapnell C et al (2022) The continuum of Drosophila embryonic development at single-cell resolution. Science 377: eabn5800

11. Campbell AG, Fessler LI, Salo T, Fessler JH (1987) Papilin: a Drosophila proteoglycan-like sulfated glycoprotein from basement membranes. J Biol Chem 262: 17605–17612

12. Cao J, Spielmann M, Qiu X, Huang X, Ibrahim DM, Hill AJ, Zhang F, Mundlos S, Christiansen L, Steemers FJ et al (2019) The single-cell transcriptional landscape of mammalian organogenesis. Nature 566: 496–502

13. Chandrasekaran V, Beckendorf SK (2003) senseless is necessary for the survival of embryonic salivary glands in Drosophila. Development 130: 4719–4728

14. Chandrasekaran V, Beckendorf SK (2005) Tec29 controls actin remodeling and endoreplication during invagination of the Drosophila embryonic salivary glands. Development 132: 3515–3524

15. Choi HMT, Schwarzkopf M, Fornace ME, Acharya A, Artavanis G, Stegmaier J, Cunha A, Pierce NA (2018) Third-generation in situ hybridization chain reaction: multiplexed, quantitative, sensitive, versatile, robust. Development 145

16. Couturier L, Mazouni K, Corson F, Schweisguth F (2019) Regulation of Notch output dynamics via specific E(spl)-HLH factors during bristle patterning in Drosophila. Nature communications 10: 3486

17. Elgar SJ, Han J, Taylor MV (2008) mef2 activity levels differentially affect gene expression during Drosophila muscle development. Proc Natl Acad Sci U S A 105: 918–923

18. Etournay R, Merkel M, Popovic M, Brandl H, Dye NA, Aigouy B, Salbreux G, Eaton S, Julicher F (2016) TissueMiner: A multiscale analysis toolkit to quantify how cellular processes create tissue dynamics. eLife 5

19. Fox RM, Hanlon CD, Andrew DJ (2010) The CrebA/Creb3-like transcription factors are major and direct regulators of secretory capacity. J Cell Biol 191: 479–492

20. Fox RM, Vaishnavi A, Maruyama R, Andrew DJ (2013) Organ-specific gene expression: the bHLH protein Sage provides tissue specificity to Drosophila FoxA. Development 140: 2160–2171

21. Friggi-Grelin F, Rabouille C, Therond P (2006) The cis-Golgi Drosophila GMAP has a role in anterograde transport and Golgi organization in vivo, similar to its mammalian ortholog in tissue culture cells. Eur J Cell Biol 85: 1155–1166

22. Girdler GC, Röper K (2014) Controlling cell shape changes during salivary gland tube formation in Drosophila. Semin Cell Dev Biol 31: 74–81

23. Haberman AS, Isaac DD, Andrew DJ (2003) Specification of cell fates within the salivary gland primordium. Dev Biol 258: 443–453

24. Hammonds AS, Bristow CA, Fisher WW, Weiszmann R, Wu S, Hartenstein V, Kellis M, Yu B, Frise E, Celniker SE (2013) Spatial expression of transcription factors in Drosophila embryonic organ development. Genome Biol 14: R140

25. Hao Y, Hao S, Andersen-Nissen E, Mauck WM, 3rd, Zheng S, Butler A, Lee MJ, Wilk AJ, Darby C, Zager M et al (2021) Integrated analysis of multimodal single-cell data. Cell 184: 3573–3587 e3529

26. Hemphala J, Uv A, Cantera R, Bray S, Samakovlis C (2003) Grainy head controls apical membrane growth and tube elongation in response to Branchless/FGF signalling. Development 130: 249–258

27. Henderson KD, Andrew DJ (2000) Regulation and function of Scr, exd, and hth in the Drosophila salivary gland. Dev Biol 217: 362–374

28. Henderson KD, Isaac DD, Andrew DJ (1999) Cell fate specification in the Drosophila salivary gland: the integration of homeotic gene function with the DPP signaling cascade. Dev Biol 205: 10–21

29. Iruela-Arispe ML, Beitel GJ (2013) Tubulogenesis. Development 140: 2851–2855

30. Isaac DD, Andrew DJ (1996) Tubulogenesis in Drosophila: a requirement for the trachealess gene product. Genes Dev 10: 103–117

31. Jacobs J, Atkins M, Davie K, Imrichova H, Romanelli L, Christiaens V, Hulselmans G, Potier D, Wouters J, Taskiran, II et al (2018) The transcription factor Grainy head primes epithelial enhancers for spatiotemporal activation by displacing nucleosomes. Nat Genet 50: 1011–1020

32. Johnson DM, Wells MB, Fox R, Lee JS, Loganathan R, Levings D, Bastien A, Slattery M, Andrew DJ (2020) CrebA increases secretory capacity through direct transcriptional regulation of the secretory machinery, a subset of secretory cargo, and other key regulators. Traffic 21: 560–577

33. Jones NA, Kuo YM, Sun YH, Beckendorf SK (1998) The Drosophila Pax gene eye gone is required for embryonic salivary duct development. Development 125: 4163–4174

34. Jürgens G, Wieschaus E, Nüsslein-Volhard C, Kluding H (1984) Mutations affecting the pattern of larval cuticle in Drosophila. Roux’s Arch Dev Biol 193: 283–295

35. Kao CF, Yu HH, He Y, Kao JC, Lee T (2012) Hierarchical deployment of factors regulating temporal fate in a diverse neuronal lineage of the Drosophila central brain. Neuron 73: 677–684

36. Karaiskos N, Wahle P, Alles J, Boltengagen A, Ayoub S, Kipar C, Kocks C, Rajewsky N, Zinzen RP (2017) The Drosophila embryo at single-cell transcriptome resolution. Science 358: 194–199

37. Kolesnikov T, Beckendorf SK (2007) 18 wheeler regulates apical constriction of salivary gland cells via the Rho-GTPase-signaling pathway. Dev Biol 307: 53–61

38. Kuo YM, Jones N, Zhou B, Panzer S, Larson V, Beckendorf SK (1996) Salivary duct determination in Drosophila: roles of the EGF receptor signalling pathway and the transcription factors fork head and trachealess. Development 122: 1909–1917

39. Lammel U, Saumweber H (2000) X-linked loci of Drosophila melanogaster causing defects in the morphology of the embryonic salivary glands. Dev Genes Evol 210: 525–535

40. Lavalou J, Mao Q, Harmansa S, Kerridge S, Lellouch AC, Philippe JM, Audebert S, Camoin L, Lecuit T (2021) Formation of polarized contractile interfaces by self-organized Toll-8/Cirl GPCR asymmetry. Dev Cell 56: 1574–1588 e1577

41. Lecuyer E, Yoshida H, Parthasarathy N, Alm C, Babak T, Cerovina T, Hughes TR, Tomancak P, Krause HM (2007) Global analysis of mRNA localization reveals a prominent role in organizing cellular architecture and function. Cell 131: 174–187

42. Levine JH, Simonds EF, Bendall SC, Davis KL, Amir el AD, Tadmor MD, Litvin O, Fienberg HG, Jager A, Zunder ER et al (2015) Data-Driven Phenotypic Dissection of AML Reveals Progenitor-like Cells that Correlate with Prognosis. Cell 162: 184–197

43. Loganathan R, Lee JS, Wells MB, Grevengoed E, Slattery M, Andrew DJ (2016) Ribbon regulates morphogenesis of the Drosophila embryonic salivary gland through transcriptional activation and repression. Dev Biol 409: 234–250

44. Lowe N, Rees JS, Roote J, Ryder E, Armean IM, Johnson G, Drummond E, Spriggs H, Drummond J, Magbanua JP et al (2014) Analysis of the expression patterns, subcellular localisations and interaction partners of Drosophila proteins using a pigP protein trap library. Development 141: 3994–4005

45. Mahaffey JW, Kaufman TC (1987) Distribution of the Sex combs reduced gene products in Drosophila melanogaster. Genetics 117: 51–60

46. Mao Y, Kerr M, Freeman M (2008) Modulation of Drosophila retinal epithelial integrity by the adhesion proteins capricious and tartan. PLoS ONE 3: e1827

47. Maruyama R, Grevengoed E, Stempniewicz P, Andrew DJ (2011) Genome-wide analysis reveals a major role in cell fate maintenance and an unexpected role in endoreduplication for the Drosophila FoxA gene Fork head. PLoS One 6: e20901

48. Maybeck V, Röper K (2009) A targeted gain-of-function screen identifies genes affecting salivary gland morphogenesis/tubulogenesis in Drosophila. Genetics 181: 543–565

49. Myat MM, Andrew DJ (2000a) Fork head prevents apoptosis and promotes cell shape change during formation of the Drosophila salivary glands. Development 127: 4217–4226

50. Myat MM, Andrew DJ (2000b) Organ shape in the Drosophila salivary gland is controlled by regulated, sequential internalization of the primordia. Development 127: 679–691

51. Myat MM, Andrew DJ (2002) Epithelial tube morphology is determined by the polarized growth and delivery of apical membrane. Cell 111: 879–891

52. Narasimha M, Uv A, Krejci A, Brown NH, Bray SJ (2008) Grainy head promotes expression of septate junction proteins and influences epithelial morphogenesis. J Cell Sci 121: 747–752

53. Nikolaidou KK, Barrett K (2004) A Rho GTPase signaling pathway is used reiteratively in epithelial folding and potentially selects the outcome of Rho activation. Curr Biol 14: 1822–1826

54. Özkan E, Carrillo RA, Eastman CL, Weiszmann R, Waghray D, Johnson KG, Zinn K, Celniker SE, Garcia KC (2013) An extracellular interactome of immunoglobulin and LRR proteins reveals receptor-ligand networks. Cell 154: 228–239

55. Panzer S, Weigel D, Beckendorf SK (1992) Organogenesis in Drosophila melanogaster: embryonic salivary gland determination is controlled by homeotic and dorsoventral patterning genes. Development 114: 49–57

56. Pare AC, Naik P, Shi J, Mirman Z, Palmquist KH, Zallen JA (2019) An LRR Receptor-Teneurin System Directs Planar Polarity at Compartment Boundaries. Dev Cell 51: 208–221 e206

57. Pare AC, Vichas A, Fincher CT, Mirman Z, Farrell DL, Mainieri A, Zallen JA (2014) A positional Toll receptor code directs convergent extension in Drosophila. Nature 515: 523–527

58. Peng D, Jackson D, Palicha B, Kernfeld E, Laughner N, Shoemaker A, Celniker SE, Loganathan R, Cahan P, Andrew DJ (2024) Organogenetic transcriptomes of the Drosophila embryo at single cell resolution. Development 151

59. Peterson J, Balogh Sivars K, Bianco A, Roper K (2023) Toll-like receptor signalling via IRAK4 affects epithelial integrity and tightness through regulation of junctional tension. Development 150

60. Sanchez-Corrales YE, Blanchard GB, Roper K (2018) Radially patterned cell behaviours during tube budding from an epithelium. eLife 7

61. Sanchez-Corrales YE, Blanchard GB, Röper K (2021) Correct regionalization of a tissue primordium is essential for coordinated morphogenesis. eLife 10

62. Schrons H, Knust E, Campos-Ortega JA (1992) The Enhancer of split complex and adjacent genes in the 96F region of Drosophila melanogaster are required for segregation of neural and epidermal progenitor cells. Genetics 132: 481–503

63. Seroka A, Lai SL, Doe CQ (2022) Transcriptional profiling from whole embryos to single neuroblast lineages in Drosophila. Dev Biol 489: 21–33

64. Seshaiah P, Miller B, Myat MM, Andrew DJ (2001) pasilla, the Drosophila homologue of the human Nova-1 and Nova-2 proteins, is required for normal secretion in the salivary gland. Dev Biol 239: 309–322

65. Sidor C, Röper K (2016) Genetic Control of Salivary Gland Tubulogenesis in Drosophila. In: Organogenetic Gene Networks, Castelli-Gair Hombría J., Bovolenta P. (eds.) pp. 125–149. Srpinger International Publishing: Switzerland

66. Street K, Risso D, Fletcher RB, Das D, Ngai J, Yosef N, Purdom E, Dudoit S (2018) Slingshot: cell lineage and pseudotime inference for single-cell transcriptomics. BMC Genomics 19: 477

67. Stuart T, Butler A, Hoffman P, Hafemeister C, Papalexi E, Mauck WM, 3rd, Hao Y, Stoeckius M, Smibert P, Satija R (2019) Comprehensive Integration of Single-Cell Data. Cell 177: 1888–1902 e1821

68. Tetley RJ, Blanchard GB, Fletcher AG, Adams RJ, Sanson B (2016) Unipolar distributions of junctional Myosin II identify cell stripe boundaries that drive cell intercalation throughout Drosophila axis extension. eLife 5

69. Tomancak P, Beaton A, Weiszmann R, Kwan E, Shu S, Lewis SE, Richards S, Ashburner M, Hartenstein V, Celniker SE et al (2002) Systematic determination of patterns of gene expression during Drosophila embryogenesis. Genome Biol 3: RESEARCH0088

70. Tomancak P, Berman BP, Beaton A, Weiszmann R, Kwan E, Hartenstein V, Celniker SE, Rubin GM (2007) Global analysis of patterns of gene expression during Drosophila embryogenesis. Genome Biol 8: R145

71. Van den Berge K, Roux de Bezieux H, Street K, Saelens W, Cannoodt R, Saeys Y, Dudoit S, Clement L (2020) Trajectory-based differential expression analysis for single-cell sequencing data. Nature communications 11: 1201

72. Wilk R, Hu J, Blotsky D, Krause HM (2016) Diverse and pervasive subcellular distributions for both coding and long noncoding RNAs. Genes Dev 30: 594–609

73. Zappia L, Oshlack A (2018) Clustering trees: a visualization for evaluating clusterings at multiple resolutions. Gigascience 7

74. Zhou B, Bagri A, Beckendorf SK (2001) Salivary gland determination in Drosophila: a salivary-specific, fork head enhancer integrates spatial pattern and allows fork head autoregulation. Dev Biol 237: 54–67

75. Zhu S, Lin S, Kao CF, Awasaki T, Chiang AS, Lee T (2006) Gradients of the Drosophila Chinmo BTB-zinc finger protein govern neuronal temporal identity. Cell 127: 409–422

76. Zhu X, Sen J, Stevens L, Goltz JS, Stein D (2005) Drosophila pipe protein activity in the ovary and the embryonic salivary gland does not require heparan sulfate glycosaminoglycans. Development 132: 3813–3822

